# Dauer quiescence as well as continuity of the life cycle after dauer-exit in *Caenorhabditis elegans* are dependent on the endoribonuclease activity of XRN-2

**DOI:** 10.1101/2022.05.02.489690

**Authors:** Tamoghna Chowdhury, Aniruddha Samajdar, Moumita Sardar, Saibal Chatterjee

**Affiliations:** Laboratory of RNA Molecular Biology Indian Institute of Science Bangalore, India

**Author notes:** Contributed equally (Tamoghna Chowdhury, Aniruddha Samajdar).

## Abstract

*Caenorhabditis elegans* embarks on a quiescent dauer state upon exposure to unfavourable conditions and can sustain for a very long period without food, but it returns to continuous life cycle upon arrival of suitable conditions. Thus, dauer state plays a critical role in its adaptive fitness and survival. ATP-independent endoribonuclease activity of XRN-2 has been implicated in dauer microRNA metabolism, perturbation of which causes their collapse within a very short span of time. Here, we present a detailed comparative analyses of dauer transcriptomes from a conditional mutant strain for the endoribonuclease activity of XRN-2, maintained under control and experimental conditions. We observed that even a limited disruption of microRNA homeostasis in experimental dauers results in deregulation of a large number of mRNA targets. Our bioinformatic analyses, supported by morphological, physiological, and behavioral evidence further demonstrate critical changes in metabolism leading to a state unsupportive of dauer maintenance, alongside potential defects in multiple neuronal activities, which might have caused an overall disruption of dauer plasticity. We explore a possible role of this endoribonuclease activity towards the maintenance of chromatin architecture and transposon expression that in turn might affect the transcriptional program critically required for the maintenance of non-aging, long-lived dauers. Finally, we also demonstrate that perturbation of the endoribonuclease activity during the dauer state exerts drastic adverse effects on the continuity of life cycle after dauer-exit. They not only fail to recapitulate the wild type events of germline development and embryogenesis, but also present traits of very old worms and formation of ‘tumor-like’ structures in the proximal gonad.

## Introduction

The quiescent dauer state in *Caenorhabditis elegans* (*C. elegans*) is a life cycle stage that has been fine-tuned through evolution to sustain unfavourable conditions such as low food availability, high population density, and high temperature^1–4^. Comparatively high *C. elegans*-specific pheromone^2, 5^ concentrations due to overpopulation, with respect to available ‘food signals’, promote dauer arrest, which further gets facilitated by high temperature. Under dauer inducing conditions, the worm first enters a ‘pre-dauer preparatory state’, L2D larva, where it senses the environmental cues through specific neurons of the nervous system and thereafter relay, process and integrate those information through dauer formation (*daf*) gene network that govern the dauer vs continuous life cycle (progression to L3 stage) developmental decision. Genetic analyses of dauer mutants have identified four distinct pathways that are involved in the regulation of dauer arrest. These pathways (Guanylyl cyclase^2, 6, 7^, TGF-β -like^2, 8, 9^, Insulin-like^2, 10, 11^, Steroid hormone pathways^12–16^) are organized in multiple layers, operate in distinct sensory and neuroendocrine tissues and form a cascade that converge on DAF-9^13, 14^, a cytochrome P450 steroid monooxygenase (3-keto-sterol-26-monooxygenase). DAF-9 executes the last catalytic step in the synthesis of a cholesterol-derived hormone in the hypodermis, somatic gonad and XXX cells, which can diffuse to most tissues to bind to the widely expressed nuclear hormone receptor DAF-12, and that in turn promotes reproductive development of the continuous life cycle^12, 15, 16^. In the absence of the hormone, DAF-12 promotes a transcriptional program towards dauer formation. A quiescent dauer worm suspends cell division, differentiation, and it is long-living as well as non-aging. However, upon arrival of favourable conditions, the dauer senses the cues and recalibrates its metabolism to get back to the reproductive phase, where it perfectly recapitulates all the molecular, physiological and behavioural programs of the continuous life cycle.

Dauer worms do not feed and hypometabolic. Their energy metabolism shifts from aerobic (TCA cycle and oxidative phosphorylation) to largely anaerobic processes, where they convert stored fats and glycogen into glycerol and glucose-1-phosphate that get metabolized through glycolysis^17–19^. Acetyl-CoA is produced from stored fat-derived long chain fatty acids through β-oxidation and that further gets converted into 4-carbon molecules (succinate, malate) via glyoxylate cycle, which are used towards the biosynthesis of carbohydrates^17–19^. Alongside glycolysis, dauer worms employ several other anaerobic pathways including malate dismutation and fermentation. Notably, malate dismutation facilitates mitochondria to function anaerobically through the use of a specialized electron transport chain^17–19^.

DAF-12, bound to its steroidal ligand, acts to oppose dauer formation, and also activates expression of *let-7* family members, which in turn suppress the expression of Hunchback-like transcription factor HBL-1^20, 21^. Otherwise, HBL-1 opposes the progression of epidermal cell fate from L2 to L3 stage. Whereas, in dauer worms, *let-7* members are diminished and that results in an increased abundance of HBL-1, which in turn favours dauer maintenance. Supported by a few more examples, thus, miRNAs are already known to be important for dauer metabolism^22^.

Recently, an XRN-2 containing macromolecular microRNA (miRNA) turnover complex (miRNasome-1) playing an important role in miRNA homeostasis has been reported from *C. elegans*^23^, where XRN-2 exhibited a previously unknown ATP-independent endoribonuclease activity^23^. Interestingly, this endoribonuclease activity was shown to be more efficacious than the previously known exoribonuclease activity, *in vitro*. Thus, it was proposed that such an ATP-independent endoribonucleolytic mode of miRNA turnover mechanism may be relevant in energy-deprived dauer worms^23^, where many miRNAs are known to be present at much lower levels compared to that of the equivalent continuous life cycle stages. Indeed, in a later study, it was found that this ATP-independent endoribonuclease activity acts on miRNAs in energy-deprived quiescent dauer worms, and its perturbation severely affects miRNA homeostasis that in turn affects the target mRNAs, and culminates in their fall^24^.

In the current study, we have performed an integrative analysis of paired miRNA-mRNA sequencing of the dauer transcriptome from a temperature sensitive mutant strain for the endoribonuclease activity of XRN-2 [*xrn-2*(*PHX25*)]. We find that impairing the endoribonuclease activity of XRN-2 by subjecting these worms to non-permissive temperature (26°C; our experimental condition, and the subjected dauers will be referred to as ‘experimental dauers’) disrupts miRNA homeostasis during dauer quiescence that in turn results in deregulation of many mRNA targets. Our bioinformatic analyses, supported by morphological, physiological, and behavioral evidence, further demonstrate signatures of gross changes in metabolism leading to a state unsupportive of dauer maintenance, alongside potentially widespread neurogenesis and neuromotor dysfunctions, and thus resulting in an overall disruption of plasticity. Moreover, we also explore a possible role of this endoribonuclease activity towards the maintenance of chromatin architecture and transposon expression that in turn might be important for the maintenance of non-aging, long-lived dauers. Finally, we also demonstrate that perturbation of the endoribonuclease activity during the dauer state exerts drastic adverse effects on the continuity of the life cycle after dauer-exit as the experimental worms not only fail to recapitulate the wild type events of germline development and embryogenesis, but also present an alarming trait of accelerated aging and formation of ‘tumor-like’ structures in the proximal gonad.

## Results

### Endoribonuclease activity of XRN-2 governs the abundance of many miRNAs in dauers that in turn modulate the levels of a large number of target mRNAs critical to multiple pathways

It has been demonstrated that the ATP-independent endoribonuclease activity of XRN-2 acts on miRNAs in energy-deprived quiescent dauer worms, and its perturbation affects miRNA homeostasis^24^. The effect of that got reflected in the deregulation of validated miRNA targets, which eventually resulted in the collapse of the sturdy dauers^24^. In the concerned study, using a low-throughput approach, the authors presented northern probing, TaqMan assays and RT-qPCR-based quantifications of a handful of miRNAs and a few of the validated targets of the cognate miRNAs that got accumulated. Although, those results were clear enough to demonstrate the role of the newly identified endoribonuclease activity in miRNA metabolism and its importance towards the maintenance of dauer stage, but it did not present a comprehensive global picture of the transcriptome indicating how the different pathways might have got affected and led to the disruption of the dauer regulatory plasticity to the extent that they collapsed within ten days. The first dauer succumbed on the fifth day (∼110 hrs) upon exposure to the non-permissive temperature, when the dauers under the control condition (20°C) remained unaffected and continued to live beyond fifty days (**Fig. 1A**). We chose this early time point (110 hrs) for performing our molecular analyses as that would exclude any secondary effect, which might get accumulated at later time points. Accordingly, the mRNA levels of miRNA biogenesis factors (eg. *drsh-1*, *dcr-1*), miRISC constituents (eg. *alg-1*, *alg-2*), and components related to miRNA turnover remained by and large unchanged (eg. *xrn-2*, *nol-58*, *miRNasome-1.4*, *paxt-1*, *xrn-1*, *dcs-1*) at that time point (data not shown).

**Fig. 1.**
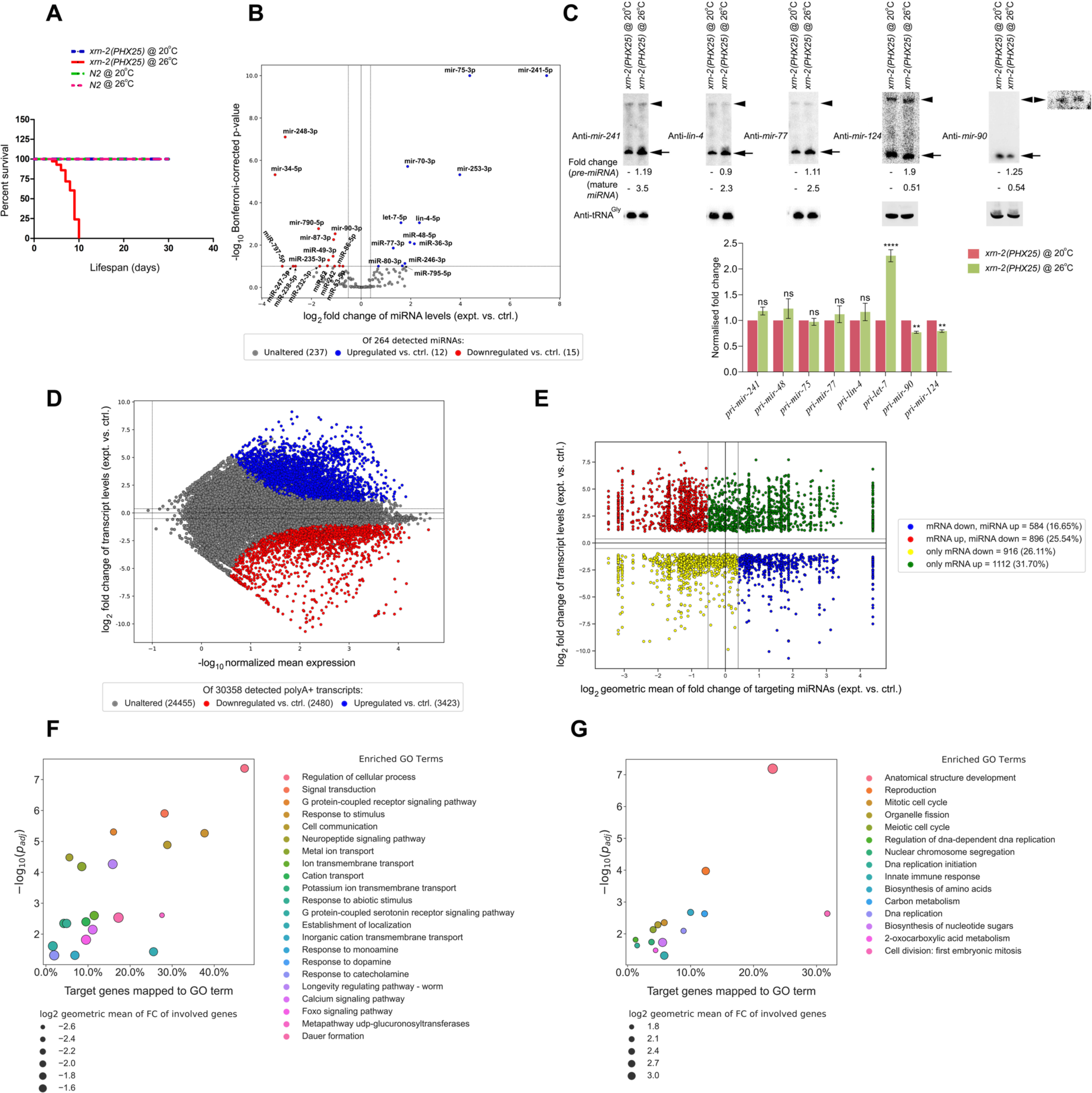
Endoribonuclease activity of XRN-2 governs the levels of many miRNAs in dauer worms that in turn modulate homeostasis of a large number of mRNAs linked to critically important biological pathways. **(A)** Dauers of the temperature sensitive *xrn-2*(*PHX25*) strain hosting mutations in the endoribonuclease active site collapse as they are subjected to non-permissive temperature (26°C). Fall of the first dauer was recorded on the fifth day (at ∼ 110 hrs), and this time point was selected for molecular analyses. **(B)** Volcano plot of expression of detected annotated miRNAs. In red are miRNAs that are downregulated at least 0.7*×* with respect to the control samples (15/264, 5.7%), and in blue are miRNAs that are upregulated at least 1.3*×* with respect to the control samples (12/264, 4.5%). Fold changes with Bonferroni-corrected p-values < 0.1 were considered significant. **(C)** Northern (**Top**) and RT-qPCR (**Bottom**) analyses demonstrate no significant changes in the levels of pre-miRNAs (indicated with arrowheads) and pri-miRNAs (except *let-7*), respectively, whose mature forms show accumulation. Two miRNAs (*mir-90, mir-124*) that are downregulated in experimental dauers (*mir-124* is not depicted in B due to *p* value cut-off) show reduced transcription and increased accumulation of their pre-miRNA forms. Top panel-extreme right depicts overexposure of a portion of the blot detecting mature *mir-90* that reveals effects on the levels of pre-*mir-90*. **(D)** MA/ Bland-Altman plot of expression of detected polyA+ transcripts for annotated genes. In red are genes whose transcripts are downregulated at least 0.7*×* with respect to the control samples (2480/30358, 8.2%), and in blue are genes whose transcripts are upregulated at least 1.3*×* with respect to the control samples (3423/30358, 11.3%). **(E)** Scatter plot of log2 fold changes in transcript levels of significantly altered genes, as described in (**D**), vs. log2 geometric mean of fold changes in significantly altered miRNAs targeting transcripts of the cognate genes, as described in (**B**). Top-left and bottom-right quadrants indicate genes whose transcript expression is inversely related with the cumulative expression of miRNAs targeting their transcripts. Top-left, in red, are gene-transcripts that are upregulated in experimental samples in correspondence with the miRNAs targeting them being cumulatively downregulated (as represented by the geometric mean), while bottom-right, in blue, are gene-transcripts that are downregulated in correspondence with the miRNAs targeting them being upregulated. **(F)** Gene Ontology (GO) Term Enrichment analysis indicates that genes downregulated with a corresponding upregulation in miRNAs targeting them under the experimental conditions are largely important for chemosensation and neural signalling. Genes related to Dauer formation and regulation of longevity also fall into this category. y-axis: significance of enrichment, x-axis: fraction of analysed genes associated with a GO term. Depicted marker radii (in black) are proportional to the geometric mean of the fold changes in expression of the genes associated with a GO term. **(G)** Gene Ontology (GO) Term Enrichment analysis indicates that genes upregulated with a corresponding downregulation in miRNAs targeting them under the experimental conditions are largely important for cell division, development (especially reproductive development), innate immune response, amino acid and other carbon metabolism. y-axis: significance of enrichment, x-axis: fraction of analysed genes associated with a GO term. Depicted marker radii (in black) are proportional to the geometric mean of the fold changes in expression of the genes associated with a GO term.

In the experimental dauers, we observed significant accumulation of 12 miRNAs, and depletion of 15 miRNAs, out of a total of 264 different species that could be detected (**Fig. 1B**). Out of 12 accumulated miRNAs, except *let-7*, all others showed no substantial changes in the levels of their pri-and pre-miRNAs (**Fig. 1C** and data not shown), and thus confirming the effect to be limited to mature miRNAs. Notably, *let-7* showed ∼2 fold upregulation in transcription (**Fig. 1C bottom panel**), whereas its mature form showed ∼3 fold accumulation compared to the control (**Fig. 1B,** a similar magnitude was also observed in TaqMan assays, data not shown). It is possible that the observed accumulation of mature *let-7* was a resultant of the events that were happening at two different levels (transcription of pri-*let-7* and degradation of mature *let-7*). Interestingly, in the experimental dauers, two of the miRNAs (*mir-90, mir-124*) that showed depleted accumulation at the level of their mature forms, also demonstrated modest downregulation of their transcripts, alongside substantial accumulation of their pre-miRNAs (**Fig. 1C)**. It indicated that not only reduced transcription, but diminished processing of their pre-miRNAs contributed to the depletion of their mature forms. This in turn suggested a possible role for the endoribonuclease activity of XRN-2 in pre-miRNA processing during a specific life cycle state, as XRN-2 depletion during the continuous life cycle is known to cause modest accumulation of the mature forms of these two miRNAs (*mir-90, mir-124*) without affecting their precursors^23^. Of note, hsXRN2 has already been implicated in the processing of pre-miR-10a in specific lung cancer cell lines (A549, H441), although, the underlying mechanistic role is yet to be uncovered^123^.

Although, only ∼4.5% of the detected miRNAs showed accumulation in the experimental dauers, but still we observed downregulation of ∼8.2% of the total transcripts that could be detected (**Fig. 1D, E**). Importantly, a large fraction of those polyA-transcripts are either validated or predicted targets having one or more miRNA binding sites in their respective 3’-UTRs^25^. Alongside, ∼5.7% of the downregulated miRNAs, we observed an increase in the levels of ∼11.3% of the detected transcripts, where a large fraction of those transcripts were validated or predicted targets of those downregulated miRNAs^25^ (**Fig. 1D, E**). Together, these extensive inverse relationships between the differentially expressed miRNAs and their cognate targets further suggested that to a reasonably large extent the effect of depletion of the endoribonuclease activity of XRN-2 on the dauer transcriptome is indeed due to its role as a ‘miRNase’, where biogenesis of the downregulated miRNAs could have been adversely affected due to depletion of transcription factors, whose mRNAs got downregulated by the accumulated miRNAs (see discussion). Thus, it appears that the endoribonuclease activity of XRN-2 directly/ indirectly affected the abundance of ∼10% of dauer miRNAs that in turn modulated ∼20% of the available polyA-transcripts, which essentially project the potential importance of this regulatory axis.

Since, the affected miRNA family members (eg. *let-7*, *lin-4* etc) are well characterised and known to be involved in the regulation of targets in different pathways, we went on to investigate the major cellular pathways that were affected in the experimental dauers. Gene Ontology (GO) terms are used to annotate genes with their functional roles in pathway databases such as the KEGG and WikiPathways databases. Therefore, we performed GO term enrichment analysis^26^ on genes, which are predicted or validated targets of affected miRNAs (as per TargetScan 7.2). GO Term Enrichment analysis^26^ indicated that genes downregulated with a corresponding upregulation in miRNAs targeting them in the experimental dauers are largely important for chemosensation and neuronal signalling, DAF-16/ FOXO pathway. Importantly, genes related to dauer formation and regulation of longevity also fell under this category (**Fig. 1F**). Conversely, GO Term Enrichment analysis^26^ indicated that genes upregulated with a corresponding downregulation in miRNAs targeting them under the experimental condition are largely important for cell division, development (especially reproductive development), as well as for the innate immune response, amino acid and other carbon metabolism (**Fig. 1G**).

### Experimental dauers show uncharacteristic downregulation of β-oxidation and glyoxylate cycle components and upregulation of TCA cycle components

Energy metabolism in dauers distinctly differs from the worms in continuous life cycle, especially the late larval stages and adults, and essentially reflects the adaptive changes indispensable for accommodating their non-feeding status^17–19^. They rely on stored glycogen^27^, trehalose^28^ and lipid^29^ for energy production. In adult worms, energy production from lipid involves β-oxidation, which converts fatty acids into acetyl-CoA, and then combustion of acetyl-CoA via the tricarboxylic acid (TCA) cycle and oxidative phosphorylation, and these combustion mechanisms are largely inoperative in dauers. In contrast to adults, in dauers, precursors for the synthesis of many biomolecules including amino acids and glucose get derived from stored fat^19, 30^. Moreover, glycerol derived from the breakdown of fat, mainly triglyceride, can be channelized into the glycolytic or gluconeogenic pathways after it gets converted into glyceraldehyde-3-phosphate, by the action of glycerol kinase and glycerol-3-phosphate dehydrogenase^31^ (**Fig. 2A**).

**Fig. 2.**
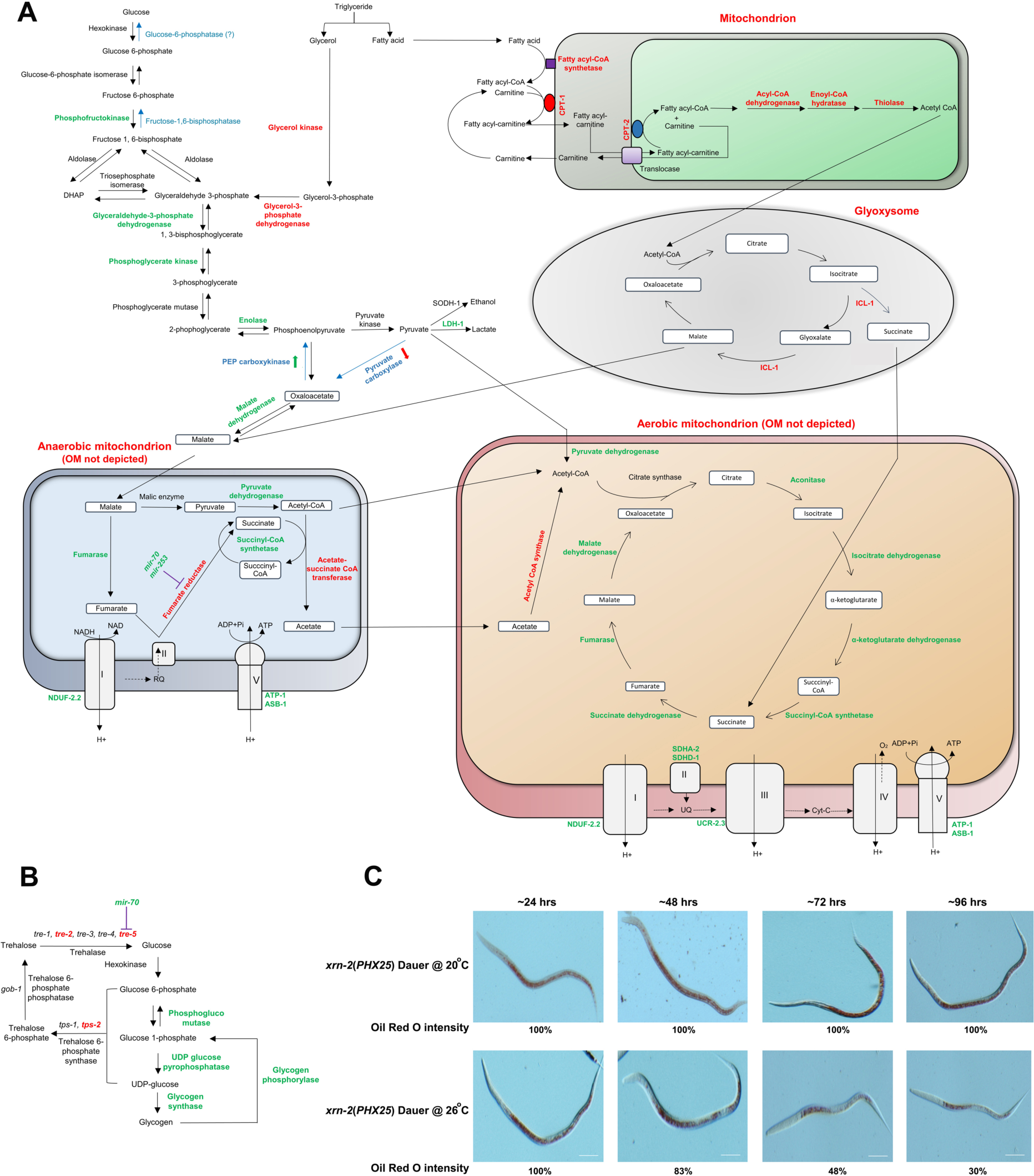
Experimental dauers show uncharacteristic downregulation of β-oxidation, glyoxylate cycle components and upregulation of TCA cycle components. **(A)** A brief overview of energy metabolism undergoing in the experimental dauers. For clarity pathways operating in their respective compartments have been depicted (mitochondrion, cytoplasm, glyoxysome). For simplicity, anerobic and aerobic mitochondria have been depicted separately. Enzymes engaged in different steps have been depicted in green or red, based on their upregulation or downregulation at the mRNA level, respectively. Critical gluconeogenic enzymes have been depicted in cyan and altered status of their mRNA levels are shown using either green-upward or red-downward arrows. A note of interrogation is used for glucose-6-phosphatase as its identity/existence is not clear. mRNA levels of enzymes driving β-oxidation and glyoxylate cycle are heavily downregulated. mRNA level of a critical enzyme of anerobic mitochondrion, fumarate reductase, shows downregulation with predicted targeting miRNAs (*mir-70*, *mir-253*) showing upregulation. Drawing has been adapted and modified from previous publications ^17, 19, 35, 36^. **(B) Trehalose pathway in *C. elegans*.** Deregulated mRNA expression of the essential enzymes of the pathway are depicted in green (upregulated) and red (downregulated). **(C)** Representative images of Oil Red O (ORO) stained *xrn-2*(*PHX25*) dauers under control and experimental conditions (20°C vs 26°C) depicting their fat reserves. Experimental dauers show dramatic reduction in ORO staining between ∼48 hrs and ∼96 hrs, following their introduction to the non-permissive temperature.

Dauers possess a functional glyoxylate pathway^32^, which allows conversion of two-carbon acetyl-CoA into four-carbon molecules that can be used as precursors for the synthesis of glucose through gluconeogenesis or formation of other cellular biomolecules^31^. Glyoxylate cycle results in the net conversion of acetyl-CoA into oxaloacetate through formation of intermediates like citrate and isocitrate, as in the citric acid cycle. Isocitrate is acted upon by the dual enzyme isocitrate lyase to produce succinate and glyoxylate, and further glyoxylate into malate. The glyoxylate cycle is completed by conversion of malate into oxaloacetate, which can be converted into phosphoenol pyruvate (PEP) by PEP carboxykinase (PEPCK). Another critical gluconeogenic enzyme is pyruvate carboxylase, which converts pyruvate into oxaloacetate (**Fig. 2A**).

In dauers, TCA cycle activities are strongly diminished relative to that of the players of glyoxylate cycle^17, 19^, which is in accordance with the prime dependence of the organism on stored fat. Additionally, it has been shown that the rate of metabolic oxygen consumption is heavily reduced in dauers^33, 34^. Interestingly, a previous study on dauers reported increased mRNA levels for genes encoding key enzymes involved in glycolysis, glyoxylate cycle, gluconeogenesis, and β-oxidation^34^. In order to understand the effect of perturbation of the endoribonuclease activity of XRN-2 on dauer energy metabolism, we compared the control dauers with experimental dauers in terms of mRNA expression of genes involved in a variety of the aforementioned key metabolic processes. Changes in mRNA levels often get transcended into changes in the abundance and thus activity of the enzymes they encode. However, one must bear in mind that the function of any enzyme is dependent on a multitude of factors like post-translational modifications, allosteric regulations, substrate availability etc.

### Effects on the players involved in glycolysis, gluconeogenesis, TCA cycle, glyoxylate cycle and fermentation

In glycolysis, glucose is converted into pyruvate, whereas in gluconeogenesis, glucose (and other biosynthetic precursors) is generated from pyruvate, as well as from other substrates such as amino acids and lactate by means of employing additional steps^31^. While most of the enzymes in glycolysis and gluconeogenesis catalyse reversible reactions, the directionality between these two processes is determined by the activity levels of a few irreversible reactions. The irreversible steps in glycolysis are catalysed by hexokinase, phosphofructokinase-1 (PFK) and pyruvate kinase (PK), whereas for gluconeogenesis, irreversible steps are carried out by phosphoenolpyruvate carboxykinase (PEPCK, although, known to act in the opposite direction under specific conditions^124^), pyruvate carboxylase, fructose-1,6-bisphosphatase (F-1,6-bisPhosphatase), and glucose-6-phosphatase (although, identity of this enzyme is not clear in worms). Dauers exhibit increased expression of hexokinase, PFK and PK, as well as the genes encoding gluconeogenic enzymes pyruvate carboxylase (*pyc-1*), PEPCK, and F-1,6-bisphosphatase (*fbp-1*). Moreover, the glyoxylate pathway enzyme ICL-1 is also upregulated in dauers. Now, compared to control dauers, the experimental dauers showed downregulation of pyruvate carboxylase (*pyc-1 mRNA*), the first critical step of gluconeogenesis. They showed no change in the level of mRNA encoding gluconeogenic irreversible enzyme F-1,6-bisphosphatase (*fbp-1*), but increased expression of the glycolytic irreversible enzyme phosphofructokinase was noted (**Fig. 2A**). Importantly, *icl-1 mRNA* was found to be downregulated in the experimental dauers (**Fig. 2A**). Collectively, the above data suggested that both gluconeogenesis and glyoxylate cycles were impaired in experimental dauers.

Pyruvate dehydrogenase (PDH) converts pyruvate to acetyl-CoA and thereby links glycolysis to the TCA cycle^31, 34^. In contrast to the scenario in control dauers, mRNA levels of the TCA cycle enzymes were dramatically upregulated in the experimental dauers, which essentially reflected an attempt by the experimental dauers to shift their energy metabolism from anaerobic to aerobic mode of operations.

In *C. elegans*, an important function of PEPCK is to drive out resources from the gluconeogenesis pathway by converting PEP into oxaloacetate. This leads to the formation of fumarate, via malate, to act as an electron acceptor in anaerobic mitochondria^31, 35^. Reduction of fumarate involves electron transfer from mitochondrial complex II to fumarate, via the electron carrier rhodoquinone (RQ) and fumarate reductase (i.e.; succinate dehydrogenase working in the opposite direction^35, 36^. It becomes possible as RQ has much lower reduction potential compared to ubiquinone (Coenzyme Q) that is integral to the succinate dehydrogenase activity driving succinate to fumarate. In another wing of this pathway, malate also gets converted into acetyl-CoA through pyruvate and forms NADH, which in turn balances the formation of NAD from NADH during the synthesis of succinate from fumarate, and thus contributes to the maintenance of cellular redox balance in the absence of oxygen.

Fumarate reductase (*F48E8.3*) catalyses the conversion of fumarate to succinate, and in dauers this gene is known to get upregulated compared to continuous life cycle worms^30, 34^. Importantly, it has been reported that the chain of reactions from PEPCK-to-succinate are critical for dauers, and thus the concerned activities get upregulated. Here, an important postulate states that anaerobic respiration involving fumarate might lower production of damaging reactive oxygen species, thereby contributing to the longevity of dauers^37^. Acetate:succinate CoA-transferase is also involved in anaerobic generation of acetate, which shows upregulation in dauer worms. Experimental dauers showed downregulation of Fumarate reductase as well as Acetate:succinate CoA-transferase (**Fig. 2A**). Notably, Fumarate reductase is a predicted target of *mir-70* and *mir-253* and both were found to be accumulated in the experimental dauers (**Fig. 2A**). Finally, changes in PEPCK levels driving oxaloacetate formation from PEP (although, there is no consensus, but a direction thought to be operational in dauers that is opposite to its normal mode of functioning, which in turn facilitates malate dismutation) would have also reduced or excluded PEP from getting channelized into the fermentation pathways, and thus in turn would have deprived the experimental dauers from accessing some important pathways of energy generation.

Trehalose, a disaccharide of glucose, is known to be a stress protectant in various species^17, 28, 32^. It is present in all the developmental stages in *C. elegans*, while most abundant in egg and dauers. Trehalose feeding extends *C. elegans* lifespan, while *daf-2*-mediated extension of lifespan is reduced by impairing genes responsible for its biogenesis, *tps-1* and *tps-2*. *C. elegans* increases endogenous trehalose production in states of extreme desiccation and during dauer state involving anhydrobiosis. The experimental dauers showed downregulation of *tps-2 mRNA*, which suggested that trehalose production might have been diminished. TRE-5^28, 32^, which converts trehalose into glucose for instant energy production in dauers and known to be dauer-enriched showed reduced mRNA expression in the experimental dauers (**Fig. 2B**). This would have affected the availability of glucose, and also might have affected gluconeogenesis, where the last step is dependent on the activity of trehalose biosynthetic pathway driving out glucose-6-p for the production of trehalose-6-p. Of note, glucose-6-phosphatase is absent or yet to be identified in worms and thus gluconeogenis is thought to be facilitated by the trehalose pathway. Overall, this data suggested that trehalose metabolism necessary for dauer maintenance might have been compromised in the experimental dauers, and thus adversely affected their survival strategies.

### Effects on mitochondrial fatty acid metabolism

It has been reported that certain β-oxidation enzymes are upregulated in dauers^34^, consistent with the fact that they are heavily dependent on catabolism of fat for energy. β-oxidation begins with the hydrolysis of triglycerides into glycerol and fatty acids, which is mediated by hormone-sensitive lipase^17^ (*hosl-1*). *hosl-1* shows upregulation in dauers, but got downregulated in the experimental dauers. Glycerol then gets converted into glyceraldehyde-3-phosphate by the action of two other enzymes namely glycerol kinase and glycerol-3-phosphate dehydrogenase. Glyceraldehyde-3-phosphate then enters the glycolytic pathway. In dauers, expression of both of these genes gets upregulated. But, in experimental dauers, these genes got downregulated at the mRNA level.

The carnitine palmitoyltransferase (CPT) shuttle system plays a rate-limiting role in the import and subsequent oxidation of fatty acyl-CoAs into mitochondria^17^. *C. elegans* genome encodes eight CPT homologs namely *cpt-1, cpt-2, cpt-3, cpt-4, cpt-5*, *cpt-6*, *b0395.3* and *w03f9.4*^17^. Majority of them shows upregulated expression in dauers. By contrast, *cpt-1, cpt-2, cpt-4, cpt-5, w03f9.4* got downregulated in the experimental dauers and among them *cpt-1, cpt-2* and *w03f9.4* are predicted to have target sites for *mir-253*, *mir-77* and *mir-246*, respectively, which showed upregulation in the experimental dauers due to compromised endoribonuclease activity of XRN-2. This might have resulted in decreased transport of fatty acyl-CoAs into mitochondria. Long-chain fatty acyl-CoA synthetases (ACS) located on the outer mitochondrial membrane (OM) regulate the rate-limiting step in β-oxidation, by activating fatty acids, through catalyzing the formation of fatty acyl-CoA. *C. elegans* genome encodes multiple potential ACS/ ligase genes, and at the least, four of them (*acs-3, acs-5, acs-17, acs-22*) show upregulation in dauers (**Fig. 2A**). In contrast to that, we observed that all these four genes were downregulated in the experimental dauers and amongst them *acs-22* has predicted target site for *mir-75*, which got upregulated in the same samples. As compared to control dauers, experimental dauers showed opposite expression patterns of the genes involved in the different steps of β-oxidation pathway (eg. components of acyl-CoA dehydrogenase, enoyl-CoA hydratase were downregulated with corresponding increase in the levels of cognate targeting miRNAs). Collectively, the above data from experimental dauers essentially reflected an overall downregulation of the β-oxidation pathway (**Fig. 2A**).

### Effects on peroxisomal fatty acid metabolism

Although oxidation of short-, medium-, long-chain fatty acids take place predominantly in mitochondria, peroxisomes are the main sites for oxidation of very long-chain fatty acids^17, 31, 38^ (C_22_ and longer). Peroxisomes also catabolize most of the less common fatty acids, including dicarboxylic acids, prostaglandins, leukotrienes and xenobiotic fatty acids^17, 31, 38^. Proteins involved in peroxisomal lipid catabolism include very long-chain acyl-CoA synthetases, ABC transporters (homologs of the adrenoleukodystrophy protein, ADLP), and peroxisomal thiolases. In dauers, majority of these genes show upregulation^39^. Moreover, peroxin genes that are involved in peroxisome assembly^38, 39^, division and inheritance are also upregulated in dauers, which essentially indicate to their elevated peroxisomal activity. Contrastingly, experimental dauers showed downregulation of all peroxin genes as well as majority of the very long-chain fatty acid transporters (*pmp-1, pmp-2, pmp-3*) and very long-chain fatty acid catabolism genes. Thus, these data suggested that experimental dauers might have suffered a compromised peroxisomal fatty acid metabolism.

Furthermore, in peroxisomes, oxidation of some fatty acids occurs slowly, and that can lead to sequestration of coenzyme A (CoASH) within peroxisomes. Acyl-CoA thioesterases act on sequestered CoASH to release fatty acyl-CoAs. In mammals, PTE-2, the broad substrate range acyl-CoA thioesterase is a key regulator of peroxisomal lipid metabolism^38^. *C. elegans* has four PTE-2 homologs, out of which three are known to get upregulated in dauers. Increased PTE activity in dauers suggests an elevated level and thus requirement for CoASH for other metabolic processes. But experimental dauers showed dramatic downregulation of all the acyl-CoA thioesterases suggesting a possible compromise in coenzyme A production.

In order to track fat utilization in control and experimental dauers, we performed Oil Red O (ORO) staining that stains fat and provides an estimation of the fat content in worms. The stained fat granules in the control dauers depicted very little change in their overall fat content from day one to day five (∼110 hrs, **Fig. 2C** and data not shown). But, in the experimental dauers, the fat content dramatically depleted between end of day two and end of day three, and further declined on day four. However, the time point (∼110 hrs, day five) at which samples were collected for RNA extraction and analyses didn’t show any significant decline in fat content from the earlier day. It is possible that fat was highly utilized between day two and day four to upregulate the expression of genes necessary for energy metabolism through TCA cycle and oxidative phosphorylation, and other necessary changes required for dauer-exit and post-dauer life (see next section). And, probably, from day four onwards there was a slow-down of energy production through fat breakdown and β-oxidation as the fat content stopped declining like the previous two days. Except one caveat described below, the mRNA expression pattern of the experimental dauers corroborated with this observation, where they showed downregulation of the genes required for dauer-specific energy metabolism and upregulation of the players involved in the energy metabolism during post-dauer stages. *lipl-5 mRNA* coding for the major fat breakdown lipase, whose expression gets stimulated upon starvation^40^, was found to be upregulated in the experimental dauers. However, this could be due to a feedback response elicited upon downregulation of β-oxidation pathway components and subsequent reduction in energy production. It could be possible that with reduced energy pool those upregulated *lipl-5 mRNAs* were not translated and thus might not have contributed to breakdown of stored fat at the given timepoint. Nevertheless, it was clear that a detailed time course study at the levels of proteins and metabolites would be necessary to comprehend the deregulated dynamics of energy metabolism in the experimental dauers that contributed to their collapse.

### Perturbation of the endoribonuclease activity of XRN-2 adversely affects the neuroendocrine pathways critical for dauer maintenance

One of the best studied examples of chemosensory seven trans-membrane spanning G-protein coupled receptors (7TM GPCRs) regulating development through cGMP signaling is the sensing of dauer pheromone, which is a measure of population density in *C. elegans*. Dauer pheromone, a complex mixture of ascarosides, is continuously secreted by *C. elegans*, and high concentrations potently induce dauer formation^5^. Several chemosensory 7TM GPCRs, including SRBC-64 and SRBC-66, SRG-36 and SRG-37, DAF-37 and DAF-38, sense specific or combinations of dauer-inducing ascarosides^41–44^. These chemosensory GPCRs signal through their G_α_ subunits, GPA-2 and GPA-3^6, 41^. When dauer pheromone levels are high, GPA-2 and GPA-3 inhibit the transmembrane guanylyl cyclase encoded by *daf-11*, thereby decreasing concentrations of the second messenger, cGMP^7^. Constitutively activated forms of GPA-2 or GPA-3, as well as loss of function mutations in *daf-11*, result in dauer constitutive (Daf-c) phenotypes, while mutations that inactivate GPA-2 or GPA-3 result in decreased dauer entry, even under dauer-inducing conditions. In our experimental dauers, genes involved in the biosynthesis of ascarosides^41^ (*daf-22*, *acox-1*, *dhs-1* and *maoc-1)* as well as genes required for the perception of dauer-inducing pheromone^5, 43^ (*daf-37*, *daf-38*, *gpa-2* and *gpa-3*) got downregulated (**Fig. 3A**). This downregulation was likely to promote the function of DAF-11^7, 9, 11, 45^, leading to a greater production of cGMP, which in turn would have promoted reproductive growth through activating TGF-β and insulin/IGF-like signaling (IIS). But *daf-11 mRNA* was also heavily downregulated in the experimental dauers (**Fig. 3A**). Therefore, despite having impaired dauer-inducing pheromone perception, the experimental dauers might have failed to produce high concentration of cGMP that is required for reproductive growth.

**Fig. 3.**
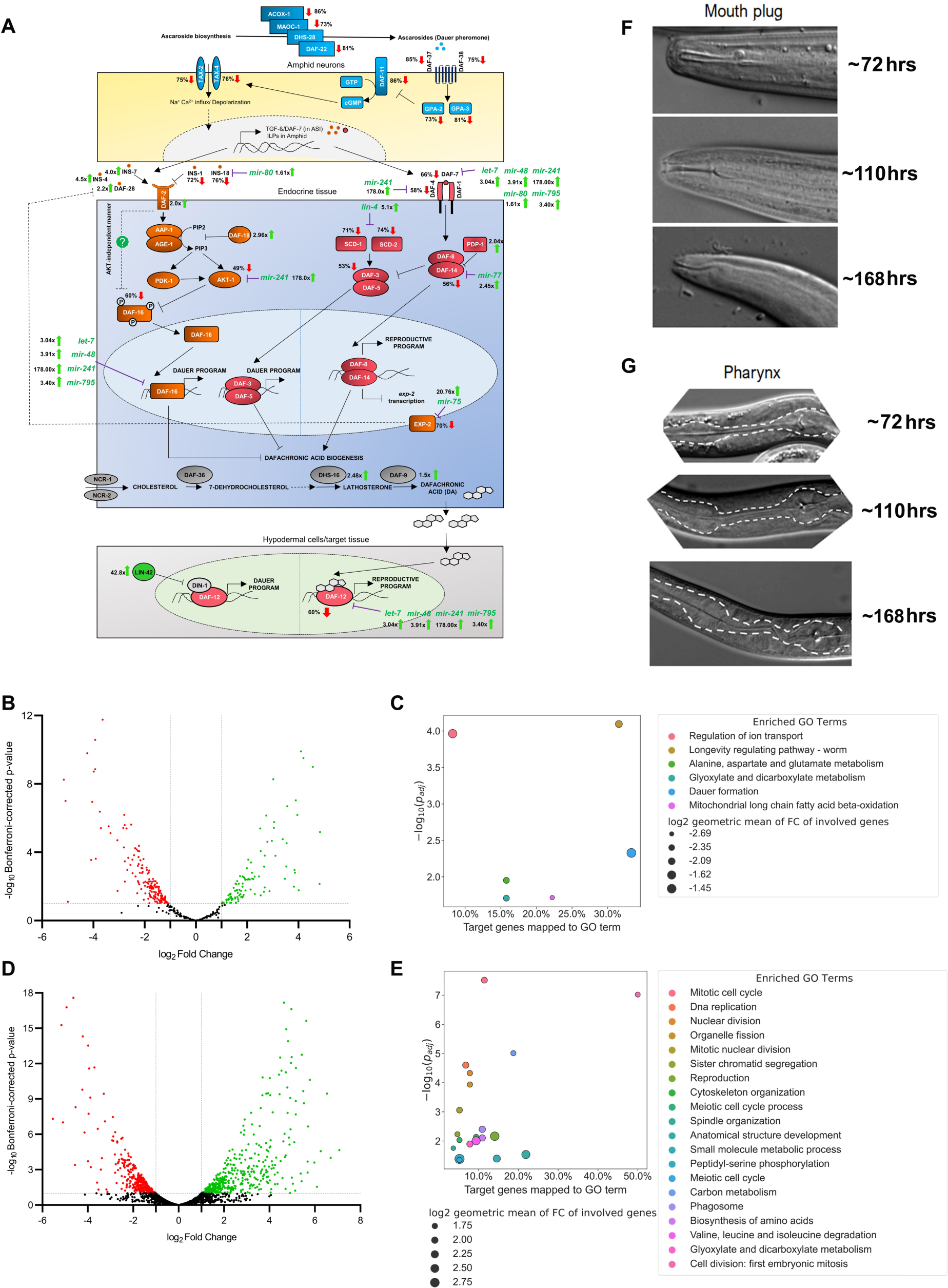
Perturbation of the endoribonuclease activity of XRN-2 adversely affects the neuroendocrine pathways required for dauer maintenance. **(A)** A graphical representation (brief overview) of the neuroendocrine axis (cGMP, IIP, TGF-β, steroid pathways) operational in experimental dauers depicting changes, compared to control dauers, in the mRNA levels of critical players, as well as changes in the levels of miRNAs targeting them. Green upward or red downward arrows indicate upregulated or downregulated mRNA levels of different components of the depicted pathways. Wherever applicable, accumulated miRNAs predicted to target the mRNAs of those components have also been depicted with green upward arrows. The endocrine pathways depict a pattern not suitable for the maintenance of the dauer state, rather facilitatory of a continuous life cycle (see text). Drawing has been adapted and modified from previous publications^13, 113^. **(B)** Volcano plot showing differentially expressed genes directly or indirectly targeted by DAF-16 (experimental vs control dauers). **(C)** Gene Ontology (GO) Term Enrichment analysis of downregulated DAF-16 target mRNAs. **(D)** Volcano plot showing differentially expressed genes directly or indirectly targeted by DAF-12 (experimental vs control dauers). **(E)** Gene Ontology (GO) Term Enrichment analysis of upregulated DAF-12 target mRNAs. **(F, G)** Experimental dauers, at indicated time points, show gradual loss of mouth plug and partial regrowth of the pharyngeal bulbs required for consumption of food.

Depending on upstream events that integrate and convey environmental cues, IIS transduces signals through a combination of finely regulated cascade of interactions^46^. During replete conditions, IIS leads to normal development and adult lifespan. Insulin-like peptides (INS-4, INS-7, DAF-28) bind and activate insulin-like receptor, DAF-2^10, 11, 29, 46^, which in turn activates AGE-1/PI3K to increase the level of PIP_3_, whose formation is antagonistically balanced by DAF-18/PTEN phosphatase by promoting the conversion of PIP_3_ to PIP_2_^47^. PIP_3_ further activates two downstream kinases, PDK-1 and AKT-1, which in turn, phosphorylates and inactivates DAF-16/FOXO transcription factor^46, 48^, by promoting its retention to cytosol with the help of FTT-2/14-3-3 proteins. Conversely, during unfavorable conditions, IIS is downregulated, allowing nuclear translocation of DAF-16/FOXO, where it turns on the expression of genes for stress resistance, dauer formation, and longevity. Notably, genetic evidence supports the existence of additional DAF-2 outputs that function in parallel to AGE-1-mediated pathway^46^. Mutations that activate *akt-1* can suppress dauer arrest in *age-1* null mutants, but they are unable to suppress dauer formation in *daf-2(e1370)* mutants efficiently. However, the weak loss-of-function allele *daf-18(e1375)* suppresses dauer arrest in *age-1* null mutants but not in *daf-2(e1370)* mutants. Furthermore, AKT-1 inactivation or mutation of consensus AKT-1 phosphorylation sites lead to DAF-16 nuclear localization, but DAF-16 is not fully active unless *daf-2* is also mutated. These results together suggest that DAF-2 exerts an inhibitory effect on the activity of nuclear DAF-16/FOXO in an AKT-independent manner.

The experimental dauers showed upregulation of *daf-2 mRNA* as well as transcripts of agonistic insulin-like peptides (INS-4, INS-7, DAF-28) and downregulation of transcripts of antagonistic insulin-like peptides (INS-1, INS-18), suggesting that the experimental dauers might have activated IIS without arriving to a condition that is favorable (**Fig. 3A**). But, these set of events probably got counteracted by upregulation of *daf-18 mRNA* as DAF-18 counters activation of PDK-1/ AKT-1 by promoting the conversion of PIP_3_ to PIP_2_. Moreover, *akt-1 mRNA*, which is predicted to have *mir-241* target site, also got downregulated. *daf-16 mRNA*, which is predicted to be targeted by *let-7* family of miRNAs, also showed 60% (*p* value = 0.03) downregulation. Therefore, it is possible that the effective protein concentrations of AKT-1 and DAF-16 must have determined the extent of phosphorylation and cytoplasmic retention of DAF-16. DAF-2, which showed a 2.0 folds upregulation at the level of mRNA, might also have influenced the activity of DAF-16 negatively in an AKT-independent manner (**Fig. 3A**). Therefore, to obtain clarity on DAF-16 functionality, we checked the mRNA levels of the genes that are known to be directly or indirectly targeted by DAF-16^48, 49^. Amongst the DAF-16 targets that were identified from mixed stage worms, 160 got downregulated in the experimental dauers (**Fig. 3B**). GO Term Enrichment analyses^26^ of those downregulated genes showed that they are indeed involved in dauer formation and longevity regulating pathways (**Fig. 3C**). This data suggested that either the concentration of DAF-16 was too low to maintain the expression of its target genes or upregulated DAF-2 via AKT-1-dependent and/or independent pathways phosphorylated majority of the available DAF-16 molecules that led to the cytoplasmic retention of DAF-16. Thus, despite food being unavailable, experimental dauers exhibited diminished DAF-16-mediated transcriptional program indispensable for dauer maintenance.

The TGF-β-like pathway that regulates dauer arrest is defined by the Daf-c genes *daf-1, 4, 7, 8*, *14* and the Daf-d genes *daf-3*, *daf-5*^50^. During replete condition, high level of secreted TGF-β family ligand DAF-7^51^ binds to and activates heteromeric cell surface receptors DAF-1^52^ and DAF-4^53^ that possess serine/threonine kinase activity, which results in the phosphorylation and activation of SMAD transcription factors DAF-8^8, 43, 54^ and DAF-14^8^. Activated DAF-8/DAF-14 complex antagonizes a complex of DAF-3/SMAD and DAF-5/SNO-SKI, which might be occurring either in the cytoplasm or nucleus, and thereby promote energy utilization and reproductive growth^50, 55^. During unfavorable conditions, reduced or absent DAF-7/TGF-β signaling permits the DAF-3/DAF-5 complex to augment dauer specific transcriptional program facilitating energy storage, and other survival traits. Experimental dauers showed reduced level of *daf-7* as well as *daf-4 mRNA* expression, in line with the fact that they are predicted targets of *let-7*s, *mir-80* and *mir-241*, respectively, which were heavily upregulated (**Fig. 3A**). Moreover, expression level of *daf-14 mRNA* was also down, as it has a predicted site for *mir-77* that showed upregulation. Accordingly, it was very likely that the TGF-β signaling pathway, promoting growth was not active in the experimental dauers, rather they might have been transducing alternative signals to maintain dauer state through the same pathway. Interestingly, the level of *daf-3 mRNA* also got diminished. Expression of two genes, *scd-1* and *scd-2*, which are thought to promote the function of DAF-3 and DAF-5 were diminished, and they are predicted to have target sites for *lin-4* that showed heavy accumulation. Thus, although, the TGF-β signaling was probably down in the experimental dauers, but it is possible that the active DAF-3/DAF-5 complex also got accumulated sub-optimally, and that might have failed to signal the worms to maintain the dauer state. Notably, DAF-8/DAF-14 activation is required for the repression of *exp-2* gene at the transcriptional level^54^. EXP-2 in turn negatively regulates the secretion of agonistic insulin-like peptide DAF-28^54^. In line with reduced *daf-14 mRNA* level in the experimental dauers, level of *exp-2 mRNA* was also much diminished and it has a predicted target site for *mir-75* that showed massive accumulation, and that in turn probably incapacitated EXP-2 to exert its inhibitory effect on DAF-28. Together, these results suggested that despite having less TGF-β signaling, the miRNA-mediated changes in the levels of some specific components might have facilitated activation of the IIS pathway to promote reproductive growth in experimental dauers.

During favorable conditions, when population density is low DAF-12 is governed by steroid hormones called dafachronic acids (DA)^13–16^. Cytochrome P450 enzyme DAF-9 catalyzes the last step (in two parallel pathways) of this hormone synthesis process^13–16^. DAF-12 acts as a developmental switch. In its DA-bound state, it promotes reproductive development, whereas in the DA-free state, it binds to DIN-1 and induces dauer formation^56^. Experimental dauers showed upregulation of mRNA levels of genes involved in the biosynthesis of DA such as DAF-9, DHS-16, suggesting that the production of DA might have been high. This presented the possibility that all the DAF-12 molecules might have been in association with DA in the experimental dauers and that in turn might have swung the decision in favour of reproductive growth. Additionally, mRNA level of *lin-42*, whose protein product blocks the DAF-12/DIN-1 complex^57^ was also dramatically high in the experimental worms, suggesting that the experimental dauers might have promoted reproductive growth by producing high amount of DA as well as by blocking DAF-12/DIN-1 complex. Notably, we also observed that the level of *daf-12 mRNA* was significantly diminished in the experimental dauers. Intriguingly, DA liganded DAF-12 promote the expression of *let-7* family of miRNAs, which in turn negatively regulate *daf-12 mRNA*, and thus form a negative feedback loop^20, 21^. Since, *let-7* family of miRNAs (*let-7*s) indeed showed dramatic upregulation in the experimental worms (**Fig. 1B, C**), due to compromised XRN-2-mediated turnover as well as DAF-12/DA-mediated transcriptional activation, they might have contributed to the downregulation of DAF-12. To better understand how DAF-12 influenced the transcriptome of the experimental dauers, we checked the expression levels of DAF-12 target genes^58^. Amongst those targets, 423 were upregulated, and 245 showed downregulation (**Fig. 3D**). GO Term Enrichment analyses^26^ of those upregulated transcripts reveled that they are involved in the reproductive growth of the organism, which otherwise must remain repressed during the dauer state (**Fig. 3E)**. These revelations supported the notion that indeed in the experimental dauers, DAF-12/DA complex was active and resulted in an altered transcriptional program that was unsupportive of dauer maintenance. In support of the above arguments, we indeed observed gradual loss of mouth plug and redevelopment of pharyngeal structures in the experimental dauers, which were indicative of their preparation for devouring food and resuming the activities of continuous life cycle (**Fig. 3F, G**).

### Experimental dauers display altered expression patterns of several genes linked with neuronal functions required for their responses to the environmental cues

In *C. elegans*, majority of the neurons are generated during embryogenesis by a series of asymmetric cell divisions oriented along the anteroposterior axis^59, 60^. Following each asymmetric division, the POP-1(Wnt)/SYS-1(β-catenin) asymmetry pathway cooperates with transcription factors (TFs) inherited from the mother cells, as well as transiently expressed TFs in some cases in the new-born precursor-neuroblast/ neuron, and collectively activate the expression of different transcription factors in the anterior and posterior daughter cells leading to the development of distinctly different neuronal cell fates^59, 60^. Combinatorial developmental inputs (lineage specific TFs and the Wnt/β-catenin asymmetry pathway) eventually initiate the expression of a unique set of TFs (terminal selectors) in each of the neuron types that leads to the expression of effector proteins necessary for the functional features/ identity of a mature neuron type, such as neurotransmitter receptors, neurotransmitter synthesizing enzymes, neuropeptides, ion channels, sensory receptors, synaptic recognition molecules etc^59–64^. These terminal selectors frequently autoregulate their expression, and they not only activate a large battery of type specific terminal differentiation genes, but also contribute to the maintenance of terminal differentiation programs in mature neuron types throughout the life of the animal. Notably, there are also pan-neuronal genes expressed in all neuron types throughout the nervous system^64^.

Dauer larvae show distinct behavioural patterns as compared to continuous life cycle worms and have altered motifs of chemosensory receptor expression^45, 65^. Results presented in the previous sections suggested that the experimental dauers showed signatures of continuous life cycle despite being famished and residing in an unfavourable environment, and that could have resulted from an incorrect assessment of the environment due to malfunctioning of the nervous system. Therefore, we were intrigued to understand whether the experimental dauers were executing the proper expression patterns of terminal selector genes, as well as pan-neuronal genes, indispensable for the proper functioning of the different neurons.

*C. elegans* hermaphrodites possess 302 neurons. White *et al.*^66^, employing highly sophisticated electron microscopic studies, accurately identified 118 classes of neurons based on their cell body positions, axodendritic features, synaptic junctions/ connectivities etc. Here, for simplicity and brevity, we aim to present how the different terminal selector gene expression patterns are affected in the experimental dauers based on the broad functional categorization of the neurons that identify them as: sensory neurons, inter-neurons and motor-neurons. Lastly, we also demonstrate the changes in the expression of genes in the experimental dauers linked to their neurotransmitter biogenesis, as well as that in the neuroglial cells.

### Sensory neurons

*C. elegans* has a powerful chemosensory apparatus that perceives chemicals in the environment, including food, noxious elements, volatile compounds, gases, mating signals, and is mediated by multiple chemosensory neurons including AWA, AWB, AWC, ASI, ASE, ASG etc^65^. Correct development/ differentiation of these neurons is essential for proper functioning of *C. elegans* sensory mechanisms. Three transcription factors namely MLS-2, CEH-36, and SOX-2 are required for AWC neuronal identity, whereas LIN-11 promotes AWA neuronal fate by inducing the expression of ODR-7, and LIM-4 along with SOX-2 controls the AWB neuronal fate^59–63, 65^. Expression of *mls-2, ceh-36, odr-7* and *lim-4* mRNAs got downregulated in the experimental dauers (the last two candidates are predicted to host *mir-80* target sites). Consequently, we observed that the direct targets (identity genes) of these terminal selectors were also downregulated in the same samples. Notably, some of these identity genes such as *odr-3*, expressed in both AWB and AWC neurons, have predicted target sites for *mir-77* and *mir-80* that were upregulated in the experimental dauers. Overall, the data suggested that the terminal selector genes as well as their targets involved in the maintenance of the differentiated state of AWA, AWB and AWC olfactory neurons, were downregulated in the experimental dauers (**Fig. 4A**). Zinc finger TF, CHE-1, along with CEH-36 drives the fate and chemotaxis properties of the ASE neurons, whereas UNC-30 plays a key role in the functionality of ASG neurons^59–63, 65^. UNC-3, a member of the Collier/Olf1/EBF (COE) TF family, induces the effector genes required for the maintenance of ASI neuron fate and functionality^59, 61, 62^. Experimental dauers showed downregulation of *che-1* and *ceh-36 mRNAs*, which might have resulted in impaired differentiation and functioning of ASE neurons. Experimental dauers also showed downregulation of *unc-3* (predicted to have target site for *mir-80*) and *unc-30 mRNAs* (predicted to have target sites for *mir-*70 and mir*-75*), required for the proper development and differentiation of ASI and ASG neurons, respectively **(Fig. 4A)**. Interestingly, *unc-30* also plays an important role in early neuronal lineage decisions as well^60^. Thus, there is a possibility that miRNA-mediated regulation of *unc-30* plays a critical role in the establishment of neuronal lineages, alongside playing important roles in the late events like terminal selection and maintenance of the differentiated state.

**Fig. 4.**
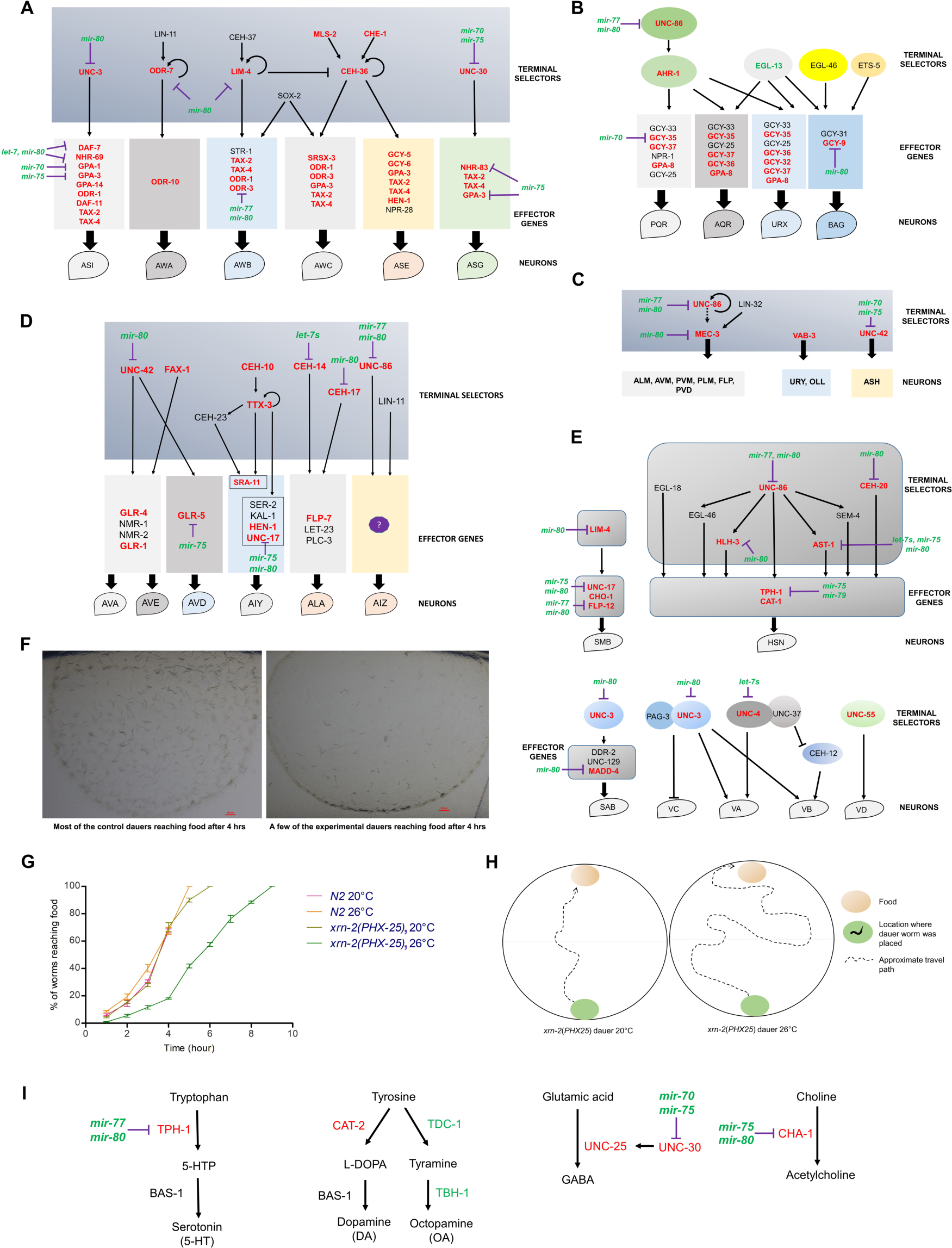
Experimental dauers exhibit deregulation of several critical mRNAs coding for terminal selectors and effectors of worm neurons that in turn are predicted targets of miRNAs showing upregulation. **(A-E)** Graphical representations (brief overview) of the developmental pathways of different neurons and the associated factors. The schemes furnish changes in the mRNA levels of critical players, as well as changes in the levels of miRNAs targeting them. Green or red coloured annotations indicate upregulated or downregulated mRNA levels of different components of the depicted pathways. Wherever applicable, accumulated miRNAs predicted to target the mRNAs of those components have also been depicted in green. Changes at many critical steps of the developmental pathways are suggestive of a pattern of deregulation that might have compromised the functioning of specialized neurons (see text). Drawings have been adapted and modified from previous publications^59–63, 65, 67, 77–79^. **(F-H)** Experimental dauers struggle to reach food, compared to control dauers. **(I)** Schematic representation of neurotransmitter biogenesis pathways. The schemes depict changes in the mRNA levels of critical players, as well as changes in the levels of miRNAs targeting them. Green or red coloured annotations indicate upregulated or downregulated mRNA levels of different components of the depicted pathways. Wherever applicable, accumulated miRNAs predicted to target the mRNAs of those components have also been depicted in green.

Additionally, *C. elegans* uses chemosensory neurons such as URX, AQR, and PQR to detect changes in O_2_ levels, thereby protecting themselves from hypoxia or hyperoxia^65^. The fates of URX, AQR, and PQR are determined by the TF, AHR-1, which in turn gets regulated by UNC-86^59–63, 65^. *unc-86 mRNA* was downregulated in experimental dauers, and it hosts predicted target sites for *mir-80* and *mir-77*, which corroboratively showed upregulation in the same samples. Notably, expression of *ahr-1 mRNA* displayed downregulation, and that in turn probably resulted in the downregulation of expression of most of its target genes, including *gcy-33*, *gcy-35*, *gcy-36*, *gcy-37* etc, which are essential for gas sensation. Moreover, some of these effector genes namely *gcy-35* and *gcy-36* have predicted target sites for *mir-7*0 that was upregulated (**Fig. 4B**). Also, *gcy-9*, a critical effector gene for BAG neurons that respond to CO_2_ stimuli, showed downregulation at the level of mRNA with a corresponding accumulation of a targeting miRNA (*mir-80*). Thus, experimental dauers might have failed to express key terminal selector genes as well as effector genes required for proper differentiation/development of neurons involved in gas sensation.

MEC-3, a LIM-homeobox TF, plays a critical role in the terminal differentiation of mechanosensory neurons such as AVM, ALMs, PVM, PLMs, FLPs, and PVDs and is required for touch response behaviour^59–63, 67^. TFs, UNC-86 and LIN-32 are the major developmental regulators of MEC-3-expressing neurons. Both factors are required for the development of the precursor lineage as well as terminal differentiation and maintenance of sensory touch neurons, and likely regulate *mec-3 mRNA* expression indirectly. In experimental dauers, both *unc-86* (hosts predicted target sites for *mir-80* and *mir-77*) and *mec-3* (hosts predicted target site for *mir-80*) got downregulated, which in turn probably led to the downregulation of downstream effector genes. Moreover, expression of *unc-42* (predicted to host target sites for *mir-70* and *mir-75*) and *vab-3 mRNAs*, which are required for the functionality of two other mechanosensory neurons namely ASH and OLL, URY, respectively, got downregulated in the experimental dauers (**Fig. 4C**).

Overall, our data suggested that genes involved in the maintenance of differentiated states of multiple neurons involved in different sensory perceptions of the worms were downregulated in experimental dauers, which might have failed them to perceive their environment correctly.

### Interneurons

After receiving environmental signals, sensory neurons relay them to the interneurons, where all the information gets processed and integrated and then transferred to downstream neurons to induce the appropriate behavioral responses. Interneurons form the main information processing/ decision-making center connecting sensory action with behavioral output.

The AIY interneurons receive and integrate information from several amphid sensory neurons, including AWA, AWB, AFD, and ASE, and are involved in locomotion, thermotaxis and chemotaxis^63, 68^. A complex comprising TTX-3 [LIM homeobox 9 (LHX9) ortholog], CEH-10 [human visual system homeobox 2 (VSX2) ortholog] along with CEH-23 control AIY neuron terminal differentiation^59–63^. These candidates are involved in an induction cascade as depicted in **Fig. 4D.** In experimental dauers, expression of *ttx-3* and *ceh-10 mRNAs* got downregulated, which might have led to compromised AIY functioning. The AIZ interneurons are involved in the odorant sensing pathway and receive signals from the AWA and AWC chemosensory neurons. UNC-86 induces AIZ generation during embryogenesis and is required to maintain AIZ function throughout life^59–63^. As mentioned earlier, *unc-86 mRNA* got downregulated in experimental dauers suggesting that AIZ neuronal maintenance might got compromised, which in turn might have resulted in defective odor-attraction and odor-adaptation behavior in them. UNC-42 plays a role in the terminal differentiation of AVD neurons, and along with FAX-1, it plays a critical role in the fate determination and axonal guidance of AVA and AVE interneurons that control backward locomotion^68, 69^. Both *unc-42* (predicted to have target site for *mir-80*) and *fax-1 mRNAs* got downregulated in the experimental dauers and that might have caused compromise in AVA, AVD, and AVE neuronal identities (**Fig. 4D**). Moreover, fate/ identity of ALA interneurons, which regulate a sleep-like behavior known as lethargus quiescence is determined by two homeodomain TFs, CEH-14, and CEH-17^70, 71^. *che-14 and che-17 mRNAs* are predicted to have target sites for *let-7* family members and *mir-80*, respectively, and displayed downregulation in the experimental dauers, which might have resulted in an improper development of ALA interneurons.

Taken together the above results suggested that terminal development and functionality of multiple interneurons might have been compromised in the experimental dauers.

### Motor neurons

Motor neurons, through the release of neurotransmitters and neuropeptides, regulate behavior-specific movements. The output of the motor neurons can be sex-specific, such that males generate mating-specific movements^63^. They can control rhythmic behaviors such as egg-laying, which is facilitated by HSN^63, 72^ (hermaphrodite-specific neuron). Along with correct fate implementation, obtainment of appropriate synaptic patterns are key for securing motor neuron diversity and function^59^.

Fate of SMB motor neuron, which influences ‘local search behavior’, is determined by LIM-4^73^. *lim-4 mRNA* has predicted target site for *mir-80* and got downregulated in experimental dauers (**Fig. 4E**). UNC-3 controls fate determination and synaptogenesis of multiple ventral nerve cord motor neurons including SAB motor neurons, which potentially control head and neck movements^74^. UNC-3 along with PAG-3 also regulates VA and VB identity, which control backward and forward movements, respectively^74^. In experimental dauers, *unc-3 mRNA* was downregulated and it has predicted target site for *mir-80*. Lack of UNC-3 in VA and VB neurons might have led to the development of an alternative VC fate in them, and that in turn might have caused defective locomotion in the experimental dauers^74^. Moreover, UNC-4 that controls the synaptic patterns in VA and VB motor neurons^74^, also got downregulated at mRNA level in the experimental dauers, which might have resulted in the generation of defective/ underdeveloped synapses with interneurons. Specific ventral and dorsal motor neurons (VD and DD motor neurons) are also involved in locomotion. The distinct synaptic patterns of these neurons are mediated by UNC-55, a nuclear hormone receptor. UNC-55 facilitates synapse formation between the VD motor neurons^75^ and the DA and DB motor neurons. *unc-55 mRNA* also showed downregulation in the experimental dauers (**Fig. 4E**), which might have led to the formation of faulty synaptic patterns leading to defective locomotion in the experimental dauers.

Finally, HSN motor neurons can control egg-laying through G protein-coupled receptor-mediated regulation of vulval muscle contraction^72, 76^. UNC-86, HLH-3, EGL-18, AST-1, SEM-4 and EGL-46 collectively control terminal differentiation and function of HSN neurons, where UNC-86 plays the role of a master regulator that influences the expression of this group of five TFs^59, 61–63, 72, 76^. Additionally, CEH-20/Pbx, another homeobox protein that does not work under the influence of UNC-86, also contributes to effector gene expression. Together, these TFs induce and maintain HSN-expressed genes, including *tph-1* and *cat-1*. Expression of *unc-86, hlh-3, ast-1* and *ceh-20 mRNAs* got downregulated in the experimental dauers. *hlh-3* and *ceh-20* are predicted to have target sites for *mir-80*, whereas *ast-1 mRNA* hosts predicted target sites for members of *let-7* family, *mir-75*, *mir-80*. *unc-86 mRNA* is also predicted to have target sites for *mir-77* and *mir-80*, and all these miRNAs were upregulated in experimental dauers (**Fig. 4E**). Interestingly, we observed egg-laying defect in these experimental worms upon dauer-exit (data not shown). Thus, upregulation of only a handful of miRNAs in the experimental dauers might have led to the downregulation of expression of some of the most important key regulator genes implicated in proper development and differentiation of neurons.

Notably, we also noticed that expression of several pan-neuronal genes such as *ric-4*, *snt-1*, *egl-3* (possesses predicted target site for *mir-36*), *rgef-1* (possesses predicted target sites for *mir-71, -75, -80*), *unc-104* (possesses predicted target sites for *mir-71, -75, -80*), *egl-21* (possesses predicted target site for *mir-80*) and *unc-10* (possesses predicted target sites for *mir-71, -80*) got downregulated in the experimental dauers, and collectively they must have affected the overall functional efficiency of the nervous system^64^.

To understand how downregulation of the aforementioned pan-neuronal, terminal selector and effector genes in the experimental dauers affected their behavior/ response to altered environmental conditions, we introduced control and experimental dauers at a given edge of petri plates and placed food (OP50) on the diametrically opposite ends of those plates. Thereafter, we monitored how much time did they take to reach the food, as well as the route and distance they traversed. Compared to control dauers, experimental dauers travelled a longer distance and took much longer time to reach the food, which indeed reflected their compromised neuronal function, although without identifying the level(s) at which the deregulations occurred to culminate into the observed outputs (**Fig. 4F-H**).

### Neurotransmitter biogenesis

Multiple genes involved in different neurotransmitter biogenesis showed altered expression levels in the experimental dauers. GABA is synthesized from the amino acid glutamic acid by the enzyme glutamic acid decarboxylase (GAD), encoded by *unc-25*^77^. Expression of *unc-25 mRNA* is dependent on the transcription factor *unc-30,* which is predicted to have target sites for *mir-70* and *mir-75. unc-30 mRNA* got downregulated in the experimental dauers, and that in turn probably led to the reduced expression of *unc-25 mRNA* **(Fig. 4I)**. Acetylcholine (ACh) is synthesized from choline by the enzyme choline acetyltransferase, CHA-1^78^. *cha-1 mRNA* is predicted to have target sites for *mir-75* and *mir-80* and was downregulated in the experimental dauers **(Fig. 4I)**.

For dopamine biosynthesis, tyrosine is first hydroxylated by tyrosine hydroxylase (CAT*-*2) to produce L-Dopa, and for 5-HT/ Serotonin synthesis, tryptophan is hydroxylated by tryptophan hydroxylase (TPH*-*1) to produce 5-hydroxytryptophan^79^. After their generation by CAT-2 and TPH-1, both L-Dopa and 5-hydroxytryptophan are then decarboxylated by the same aromatic amino acid decarboxylase (AAAD), encoded by *bas-1*, to produce dopamine and serotonin, respectively^79^. Whereas, octopamine biosynthesis depends on tyrosine decarboxylase (TDC-1), which converts tyrosine into tyramine. Then tyramine is converted to octopamine by tyramine β-hydroxylase^79^. Both *cat-2* and *tph-1 mRNAs* got downregulated in the experimental dauers (**Fig. 4I**), suggesting a possible compromise in the production of both dopamine and serotonin. Interestingly, mRNA levels of both octopamine biosynthetic enzymes showed upregulation in the experimental dauers. In the next section, we will discuss the downregulation of the octopamine receptor (OCTR-1) at the level of its transcript. Thus, it is possible that increase in the mRNA levels of octopamine biosynthesis components might have happened because of feedback response for downregulated *octr-1* expression.

Lastly, **glial cells** are known to be important for the homeostasis of the mammalian differentiated neurons and recovery of post-injury neuronal tissue, and thus for better understanding of the subject, worm glial cells are also actively investigated^80, 81^. CEPsh glial cells in worms have been shown to modulate dopamine-dependent behaviors, including feeding and other learning behaviors^81^. Disruption of the *hlh-17* gene led to changes in egg-laying behavior, feeding behavior-plasticity deficits, and an impaired form of gustatory associative learning^81^. The four CEPsh glial cells present in worms are closely associated with the four CEP neurons, which facilitate the aforementioned behaviors through release and up-take of dopamine. Developing CEPsh glia are known to specifically express HLH-17, an ortholog of the TF, OLIG2, which is required for oligodendrocyte specification^80^. In the developing spinal cord of vertebrates, oligodendrocytes form distinct ventral and dorsal domains, where *Olig2* expression is induced in the ventral region by the TFs, Nkx6 and Pax6. Interestingly, CEPsh expression of *hlh-17* also requires the Nkx-like *mls-2* and Pax-like *vab-3*^80^ (*mir-235*, *mir-249*, *mir-63*s, and *mir-231*, *mir-787*, *mir-236*, *mir-50*s are predicted to target *mls-2* and *vab-3 mRNA*s, respectively). Now, all these three genes showed dramatic downregulation in the experimental dauers (data not presented), and that might have affected them adversely. Of note, we did not observe any changes in the levels of the miRNAs potentially targeting those downregulated candidates. Thus, it would be necessary to perform an in-depth systematic study to understand the role of XRN-2-mediated regulation of miRNAs, and in turn their direct and indirect target mRNAs, in worm glial cells, if any, and how that affects the modulatory role they play on other neurons.

### Immune system components of the experimental dauers exhibit altered expression patterns

The worm innate immune system provides the first line of defence against pathogenic infections and relies upon pathways that are conserved across phyla. It is known that the worm nervous system elicits and regulates the innate immune response upon exposure to pathogens, and neuronal GPCRs play critical roles in inducing and governing several key downstream pathways that mount the immune response^82, 83^. Since, in the earlier section, it could be noted that neuronal development and functions potentially get shaped by miRNAs, which in turn get regulated by the endoribonuclease activity of XRN-2, we were intrigued to understand the overall changes in the expression of the components of the immune system that might have occurred in the experimental dauers.

In *C. elegans,* evolutionarily conserved p38 mitogen-activated protein kinase cassette comprising PMK-1, NSY-1, SEK-1, TIR-1 plays a central role in immune response, where PMK-1^84, 85^ is the most well studied effector protein (p38 MAPK PMK-1). This pathway governs the infection-induced expression of secreted immune response gene products, including C-type lectins, lysozymes, and antimicrobial peptides that are known to fight off infection in many species. However, host innate immune responses need to be tightly regulated because insufficient response may exacerbate infection, whereas excessive response may result in prolonged inflammation, which might lead to tissue damage and death. Therefore, multiple pathways exist that under normal conditions suppress the unwanted activation of p38 MAPK pathway, and thereby protect the organism from unwarranted activation of innate immune effectors.

*C. elegans* operates multiple immuno-suppressive pathways mediated by receptors OCTR-1^83^, OLRN-1^82^ and dopamine-DOP-4^86^. Neurotransmitter octopamine (OA) released from RIC interneurons, acts as a ligand for OCTR-1 that functions in the sensory ASH neurons, which leads to the suppression of innate immune response in pharyngeal and intestinal tissues^83^. Ligand activated OCTR-1 suppresses the expression of non-canonical unfolded protein response (UPR) genes of the *pqn/abu* family, as well as genes in the immune pathway mediated by p38 MAPK PMK-1. Notably, OA is the invertebrate counterpart of the vertebrate neurotransmitter norepinephrine. The OA receptor, OCTR-1, is orthologous to the alpha-2A adrenergic receptor of norepinephrine. Since, norepinephrine regulates innate immune responses in mammals and other vertebrates, the immunomodulatory functions of OA/norepinephrine are thought to be conserved. Sympathetic noradrenergic neurons, especially splenic nerve, function in the mammalian inflammatory reflex to suppress infection-induced production of inflammatory cytokines in the spleen and other organs. Thus, this immune-inhibitory neural circuit plays a critical role in maintaining optimal immunogenic response. Although, genes involved in the biogenesis of OA got dramatically upregulated in the experimental dauers, *octr-1 mRNA* (predicted to have target site for *mir-75*) expression was downregulated. Consequently, it could be noted that the mRNAs of non-canonical UPR genes of the *pqn/abu* family and effector genes of the p38 MAPK PMK-1 immune pathway also got dramatically upregulated (**Fig. 5A**), and that too in the absence of any pathogen.

**Fig. 5.**
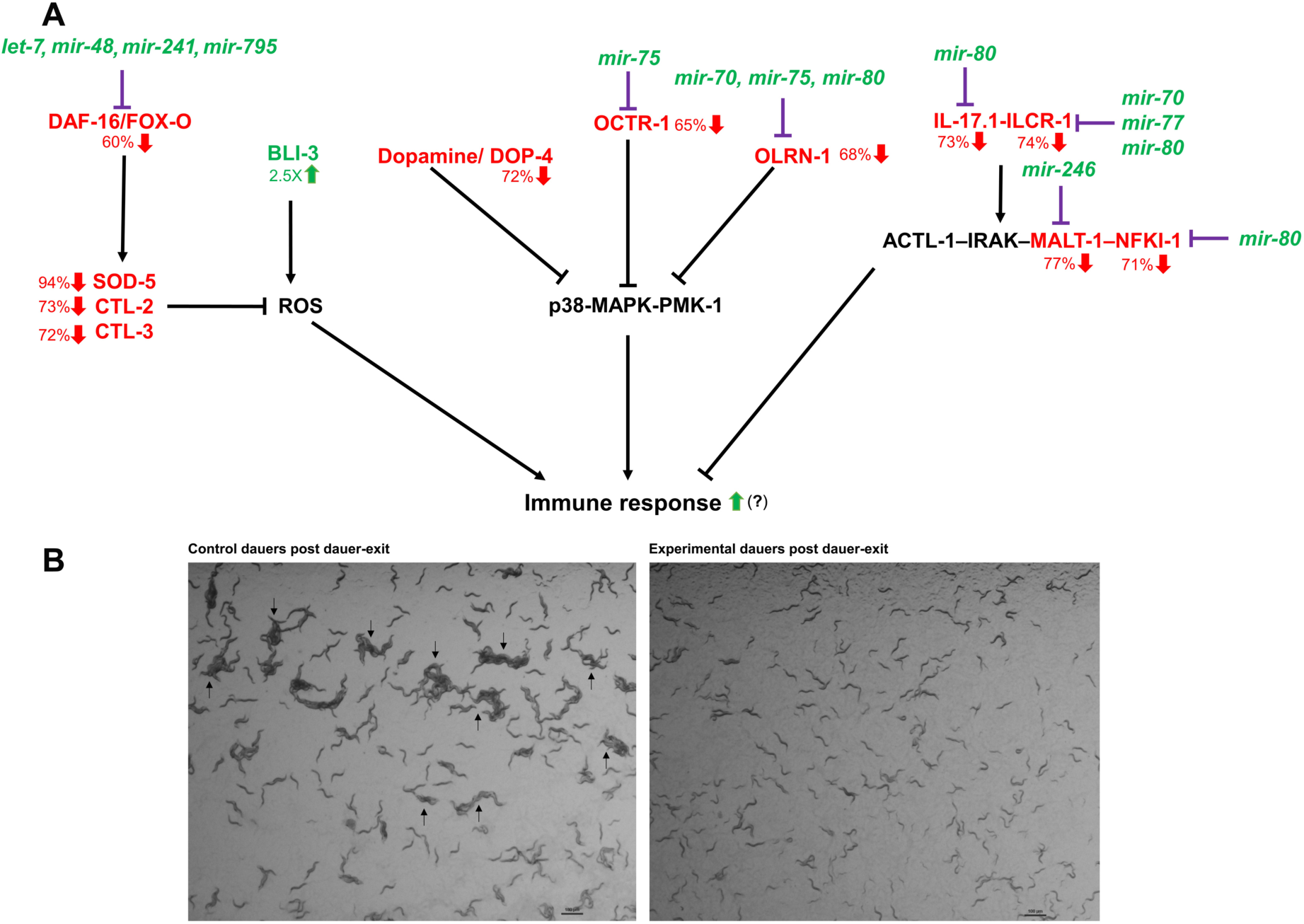
**(A) Experimental dauers exhibit altered expression of immune system components.** Changes, compared to control dauers, in the mRNA levels of critical players, as well as changes in the levels of miRNAs targeting them have been depicted through green (upregulation) or red (downregulation) coloured annotations. **(B) Experimental dauers (right panel) show reduced aggregation property** depicting inability to perceive high oxygen and perform aversion behaviour that is linked to reduced expression of IL-17 pathway components (see text, aggregates are indicated with arrows).

Another GPCR, OLRN-1, functions in AWC neurons in the cell non-autonomous suppression of the canonical p38 MAPK PMK-1 immune pathway in the intestine^82^. *olrn-1 mRNA* is predicted to have target sites for *mir-70, mir-75* and *mir-80* and showed dramatic downregulation in the experimental dauers. Moreover, dopamine, through DOP-4, a D1-like dopamine receptor, suppresses innate immune responses in worms by downregulating the p38 MAPK PMK-1 pathway. Genes involved in dopamine biogenesis as well as *dop-4 mRNA* were heavily downregulated in the experimental dauers (**Fig. 5A**).

In light of the above observations, it appeared that the pathways that are responsible for immune suppression were downregulated in the experimental dauers. This is in corroboration with the finding that expression of genes involved in the immune response, including production of C-type lectins, lysozymes, lipases and antimicrobial peptides, were heavily upregulated in the experimental dauers (data not shown). As excessive innate immune responses have been linked to human health conditions such as Crohn’s disease, rheumatoid arthritis, atherosclerosis, diabetes, and Alzheimer’s disease^87^, XRN-2’s endoribonuclease activity projects a potential to play a key role in the regulation of innate immunity and auto-immunity through its ability to modulate miRNA landscape under specific conditions.

In addition to its arsenal of antimicrobial proteins and peptides, *C. elegans* also has the capacity to produce bactericidal reactive oxygen species (ROS) in response to exposure to pathogens via the action of the dual oxidase, BLI-3^88^. But, ROS do not act on specific targets, they indeed damage host tissue alongside killing pathogenic bacteria. As a compensating mechanism, increased levels of ROS trigger a protective stress response in the host involving upregulation of the members of the superoxide dismutase family (eg. SOD-1/2/3/4/5) and catalases (eg. CTL-1/2/3), whose production are known to be dependent on DAF-16 activity (see earlier results section). Even in the likely absence of any bacterial pathogen in the immediate environment, the experimental dauers showed upregulation of *bli-3 mRNA*, but *ctl-1*, *ctl-2* (predicted to have target site for *mir-75*), *ctl-3 mRNAs* as well as *sod-5 mRNA* got heavily downregulated and that is probably due to reduced DAF-16 activity. Therefore, it could be possible that ROS produced by BLI-3 resulted in tissue damage in the experimental dauers and contributed to their collapse.

Interleukin-17 (IL-17) is a pro-inflammatory cytokine^89–91^ and its mRNA level showed diminishment in the experimental dauers. In mammals, it is involved both in host defence and in pathological inflammatory conditions such as autoimmune diseases. IL-17 is produced predominantly by the Th17 cells, whose differentiation from precursor T cells depend on IL-6 and TGF-β. Again, it is to be noted that in quiescent dauers the expression of the latter candidate (DAF-7/TGF-β) gets modulated by XRN-2 regulated miRNAs. IL-17 facilitates numerous immune and inflammatory responses by regulating the expression of various inflammatory mediators, which include cytokines, chemokines, and adhesion molecules. It is involved in the pathogenesis of asthma, where it orchestrates the neutrophilic influx into the airways and enhances T-helper 2 (Th2) cell-mediated eosinophilic airway inflammation in mice, and similar events have also been recorded in patient samples^91–93^. IL-17 has also been implicated in promoting inflammatory response and inducing secondary injuries to post-stroke tissue. In *C. elegans*, rather surprisingly, neuronal production of IL-17 suppresses intestinal defence mechanisms against infection^89, 90^. It has also been shown that IL-17 and its receptors ILCR-1/ ILCR-2, as well as the downstream ACTL-1–IRAK–MALT-1–NFKI-1 complex-mediated signalling are required in one hub interneuron to increase its response to presynaptic input from oxygen sensors. This allows *C. elegans* to persistently escape high oxygen concentrations. Thus, IL-17 signalling promotes aversion to noxious stimuli. IL-17 neuronal activity allows integration of sensory inputs and appropriate cognitive behaviour in multiple species. Coordination of the immune response with an appropriate behaviour (escape, learning, social interactions) likely increases organism’s fitness. Now, the mRNAs of *ilc-17.1* (IL-17) and its receptor *ilcr-1* have target sites for *mir-80* and *mir-70*, *mir-77*, *mir-80*, respectively. Whereas, *nfki-1* and *malt-1 mRNAs*, downstream components of this pathway, host conserved target sites for *mir-80* and *mir-246,* respectively. All these miRNAs were upregulated in the experimental dauers and accordingly, expression of *ilc-17.1*, *ilcr-1*, *nfki-1* and *malt-1 mRNAs* were downregulated in the same sample. Importantly, the experimental dauers, upon introduction to a seeded plate, were unable to aggregate to preclude themselves from high (21%) O_2_ (**Fig. 5B**) and that indeed was a manifestation of impaired signalling events linked to O_2_ aversion behaviour.

### Endoribonuclease activity of XRN-2 affects transposon metabolism and the components of chromatin dynamics in dauers

Wild type *N2* worms can live up to ∼17 days^94^, but dauers can sustain up to 3-4 months without any food intake, remaining dependent on their internal energy reserves. Although, dauers are metabolically much less active compared to worms in the reproductive phase, but a specialized metabolism allows them to sustain the adverse conditions. Accordingly, dauers adopt a transcription program, distinctly different in many aspects from the continuously growing worms^30, 95^. Thus, it is very much possible that specialized chromatin architecture and specific epigenetic modifications might allow the dauers to maintain a unique transcriptional program for a duration that is ∼ten times longer than its normal life span. Therefore, we were intrigued to know whether specific epigenetic modifiers and chromatin components exhibited altered expression in the experimental dauers that failed to sustain beyond ten days.

We observed that MES-4, a germline enriched H3K36 histone methyl transferase^96, 97^, got upregulated in the experimental dauers. In *C. elegans*, H3K36me3 and H3K27me3 occupy unique loci on autosomes, and methylation of H3K36 adversely affects H3K27me3 marks placed by Polycomb Repressive Complex 2^97^ (PRC2, comprising MES-2/3/6, **Fig. 6A**). Notably, PRC2 mediated formation of H3K27me3 marks and MES-4 mediated H3K36me3 marks counterbalance each other to ensure correct gene expression in the germline. MES-4 function alleviates the germline specific autosomal genes from the repressive activity of PRC2 and limits its repressive action on other autosomal regions harbouring somatic genes, as well as on genes housed in X chromosomes. MES-4 upregulation in experimental dauers might have led to changes in the *bona fide* distribution of H3K36me3 and H3K27me3 patterns in the genome and resulted in the expression of those germline enriched genes (**Fig. 6A**), which otherwise must have remained repressed during dauer quiescence. In support of this notion, we also observed that the germline genes that remain associated with MES-4^96^, showed upregulation in the experimental dauers (**Fig. 6B**).

**Fig. 6.**
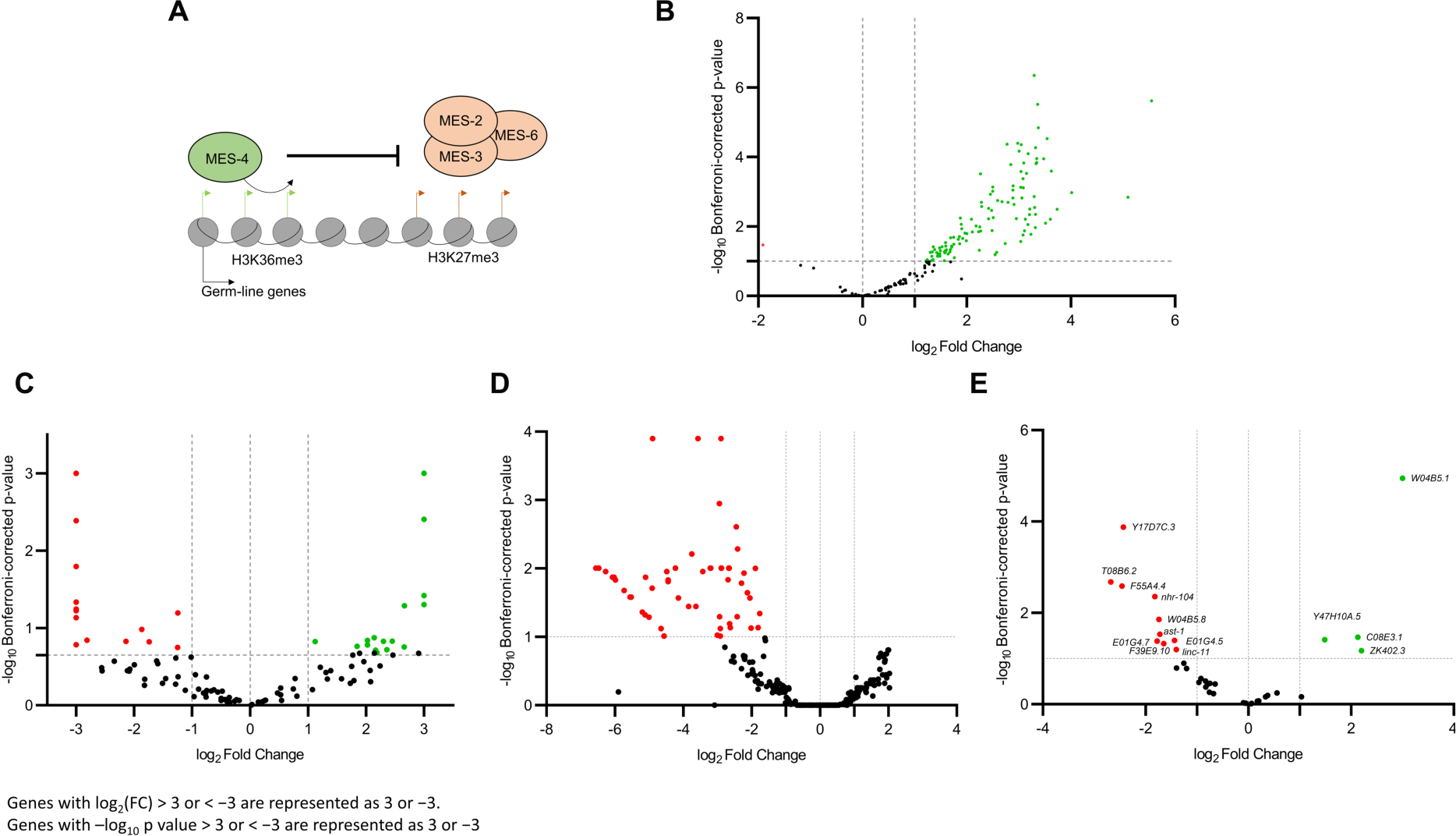
Experimental dauers show deregulation of players important for chromatin dynamics, as well as altered endo-siRNA and transposable element metabolism. **(A)** Schematic representation of *mes-4* upregulation (depicted inside a green oval) leading to altered chromatin architecture/ dynamics. **(B)** Volcano plot showing differentially expressed germline genes that remain associated with MES-4. **(C)** Volcano plot showing differentially expressed transposable elements in experimental dauers compared to control dauers. **(D)** Volcano plot showing differentially expressed endo-siRNAs/ 22G-RNAs in experimental dauers with respect to control dauers. **(E)** Volcano plot showing differentially expressed mRNAs that got targeted by the differentially accumulated 22G-RNAs in experimental dauers in comparison to control dauers.

We noted that the *C. elegans* orthologue of JMJD3, JMJD-3.1, which plays an important role in H3K27me3 demethylation^98^, got downregulated. Notably, this protein is known to be essential for proper gonadal development^98^. Accordingly, after dauer-exit of the experimental dauer worms, upon exposure to food, a plethora of defects in gonadal arm development could be recorded (see next section).

In dauers, expression of replicative histones remains silenced. Only two histone H1-like genes, *hil-1* and *hil-3*, show increased expression and have been implicated in dauer longevity^95^. Both *hil-1* and *hil-3* host predicted target sites for *mir-80* and were downregulated in the experimental dauers, whereas several other replicative histones showed upregulation (data not presented). Collectively, these might have resulted in altered chromatin architecture/ dynamics not conducive for the gene expression program of dauers.

In all organisms, presence of repetitive and mobile elements within the genome present grave danger to the integrity of the genome and fidelity of the genetic information, which would eventually compromise the overall fitness of the organism. Accordingly, molecular machinery evolved that suppress repeat sequence expression and transposable element (TE) mobility within the genome. In *C. elegans*, WAGO/22G-RNAs play a critical role in silencing of TEs^99, 100^. Previously, XRN-2 was implicated in the silencing of several classes of TEs, where upon depletion of XRN-2, 22G-RNAs targeting different TEs got dramatically downregulated^101^. Thus, we asked whether TEs got derepressed in the experimental dauers, and indeed we identified sixteen different TEs, including *cer17-i*, *tc1a*, *mirage1*, WBT00000684, WBT00000756, WBT00000717 and WBT00000759, which showed upregulated expression in experimental dauers, whereas fifteen showed downregulation (**Fig. 6C**). Interestingly, in case of *mirage1*, we could also note downregulation of corresponding 22G-RNAs, although with low significance. Probably, with a higher depth and coverage in the small RNA sequencing data, for rest of the TEs, one would be able to detect the corresponding 22G-RNAs. Now, in worms, it is not clear whether majority of the TEs do have their own promoters or do they largely depend on the promoters of the neighbouring genes and host genes for their expression^125^. Of note, in mammalian systems, extensive work has shown that TF binding sites embedded within TEs indeed have shaped the gene regulatory networks of the hosts not only by regulating transcription, but also through modulating genomic organization, especially in germ cells and embryos during a specific time-window (pre-implantation development)^126, 127^. Since, we have seen altered expression of a large number of genes in experimental dauers, it is also possible that the changes in the transposase mRNA levels were fully/ partly due to the differential expression of their host or neighbouring genes. Also, many mammalian TFs have been shown to perform sequence specific binding of TEs resulting in their expression^126, 127^ and some of the worm homologs of such TFs (eg. *sox-4*, *rnt-1*, homologs of mammalian sox11, runx3, respectively) were found to be showing deregulated expression in the experimental dauers, which could have potentially affected the expression and thus mobilization of the concerned TEs, alongside affecting the expression of the adjacent and/or host genes. Analyses of CHIP-seq experimental data^128^ for *sox-4* and *rnt-1* might reveal the identity of the TE families to which they bind (**Fig. 6**), alongside other growth-promoting genes and some of those might also come under the sphere of influence of the newly mobilized TEs. Thus, a detailed CHIP-seq analyses of the different critical TFs in the experimental dauers, along with an understanding of the physical locations of the concerned TEs, would be necessary to understand how XRN-2 potentially affects the dynamics of interactions of TEs and TFs and their effects on different networks of gene expression. Interestingly, we observed that mRNAs coding for proteins having roles in non-homologous end-joining (e.g. *mre-11* and *cku-70*) got upregulated in the experimental dauers. Mobilization of DNA cut-and-paste TEs (eg. *tc1a*, *mirage1* etc) leads to dsDNA breaks, and in response to that non-homologous end-joining pathway gets activated. Moreover, genes coding for multiple DNA repair pathway components such as *msh-6, msh-2, xpb-1, xpc-1, exo-3, brc-1* (the worm homolog of mammalian BRCA-1) etc. got upregulated in the experimental dauers. Collectively, these results were suggestive of the notion that the experimental dauers experienced increased genotoxic stress and transposon mobilization might have occurred, which ultimately adversely affected their genomic stability and might have also contributed to the development of germline ‘tumor-like’ structures post dauer-exit (see next section). Data from future genome sequencing experiments would help us to achieve resolution on this notion.

It has already been demonstrated that CSR-1 pathway positively influences the expression of genes involved in the detection of different environmental stress conditions by the sensory neurons, as well as many of the associated signalling events^102^. Thus, it is possible that CSR-1 plays an important role in dauer formation and its subsequent maintenance. Recently, we have shown that in the continuous life cycle, XRN-2 is essential for the accumulation of different types of endo-siRNAs. Therefore, we were curious to know whether endo-siRNAs displayed differential expression in the experimental dauers. Overall, we could detect endo-siRNAs (22G-RNAs, **Fig. 6D**) with confidence against 1070 transcripts, and amongst them 74 displayed deregulation of expression compared to the control dauers. Those 74 transcripts code for 48 unique genes. Out of those 48 genes, ten showed dramatic downregulation, whereas four were upregulated at the mRNA level (**Fig. 6E**). Interestingly, majority of those downregulated genes are known to be neuron specific and get heavily expressed during dauer life cycle stage, but they are largely uncharacterized. Therefore, it will be interesting to study the roles that those genes might play in the maintenance of dauers.

### Deregulated miRNA homeostasis during dauer state adversely affects events of continuous life cycle after dauer-exit

It has already been shown that post-dauer worms display altered chromatin marks as well as altered gene expression patterns, in addition to expression of genes involved in adult life-span regulation and reproduction, compared to the worms that have been growing continuously^103^. Consequently, the post-dauer worms produce more progeny and live (healthily) longer than those continuously growing worms that would show signs of senescence after a given number of generations. In a nutshell, dauer quiescence facilitates rejuvenation of the immortal germline phenotype upon dauer-exit.

The experimental dauers in our experiments, when provided with food, showed a plethora of developmental defects upon dauer exit. Only 62.5% (± 2.9%) reached the adult stage (**Fig. 7A left panel**), but they displayed multiple defects in gonadal development and traits normally seen in very old worms **(Fig. 7B, C,** data not shown). We not only observed morphological defects in the developing oocytes **(Fig. 7Biii, 7Cii, Ciii)**, but also noted their maturation in ectopic locations, especially towards the distal part of the gonadal arms **(Fig. 7Biv)**. We also observed that 27% (±2.7%) of the total worms or 45% (± 7%) of the worms that reached adult stage showed ‘tumor-like’ growth near the proximal ends of their gonadal arms (**7Biii-vi**), and that was reminiscent of germline teratomas normally observed in very old worms. Those worms also displayed defects in embryogenesis (**Fig. 7D,** data not shown), and had an average brood size of 20 (± 2) upon return to the continuous life cycle, but became completely sterile and ceased to continue the life cycle within three generations post-dauer-exit (**Fig. 7A right panel**). Conversely, the *xrn-2(PHX25)* control dauers (maintained at 20°C) as well as *N2* dauers maintained at 20°C or 26°C, when provided with food, embarked on the reproductive cycle of life, and displayed normal development for the subsequent many generations **(Fig. 7A** and data not shown).

**Fig. 7.**
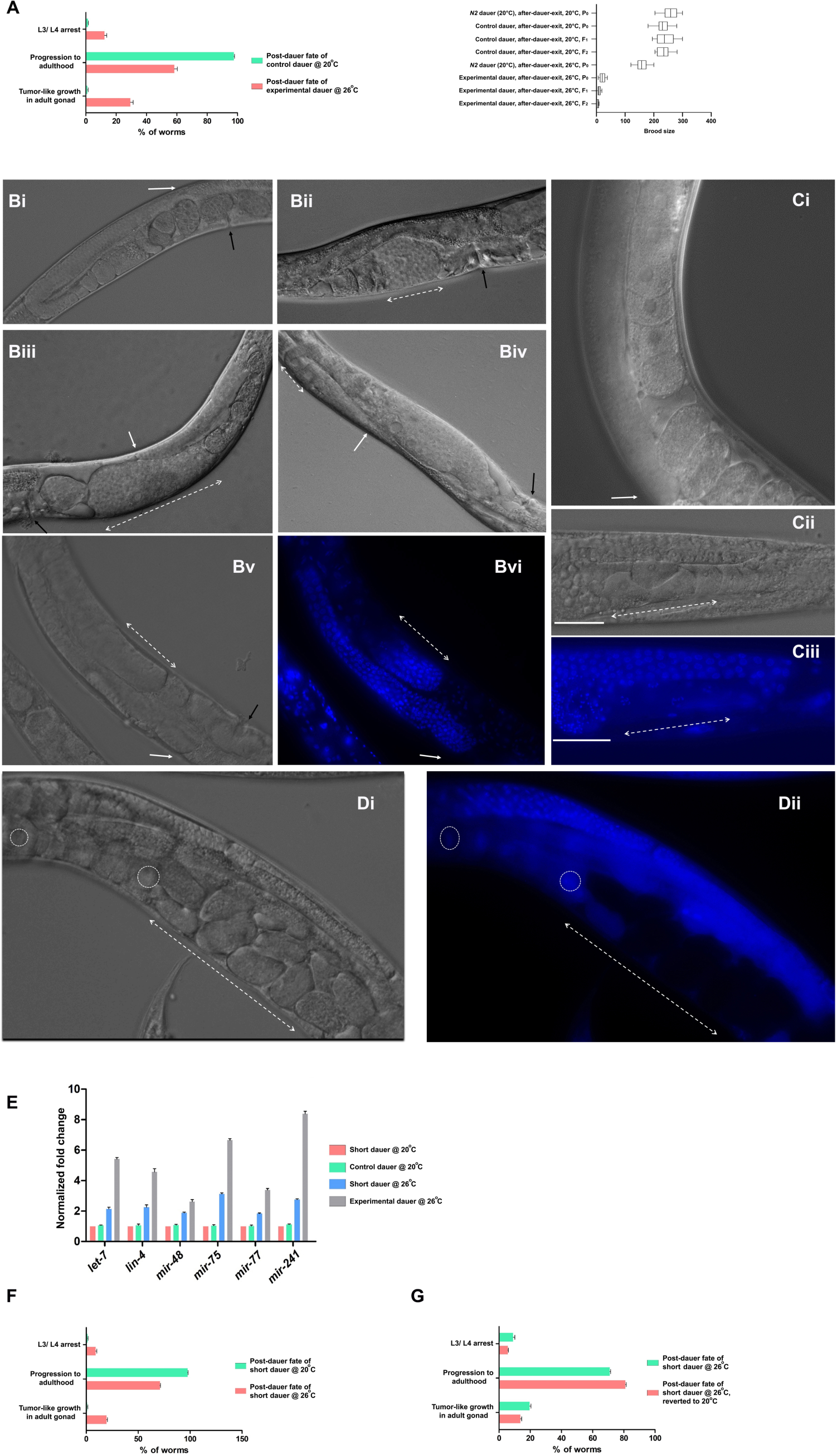
Deregulated miRNA homeostasis during dauer state adversely affects events after dauer-exit. **(A) Left panel.** Experimental dauers perform dauer-exit after introduction to food but show defects during larval development and after reaching adulthood, as indicated. **Right panel.** Box plot showing brood size of different post-dauer worms, as indicated. **(Bi)** A representative DIC image of the adult stage of a control dauer [*xrn-2*(*PHX25*)] after dauer-exit at permissive temperature (20°C). White arrow indicates to the distal tip of the gonadal arm, while the black arrow indicates to vulva. **(Bii)** A representative DIC image of young adult stage of an experimental dauer [*xrn-2*(*PHX25*)] after dauer-exit at non-permissive temperature (26°C). Black arrow indicates to the position of vulva. Double-headed arrow indicates to a ‘nuclei mass’ located between ‘malformed’ oocytes and vulva. **(Biii)** Early gravid adult stage of an experimental dauer [*xrn-2*(*PHX25*)] after dauer-exit at non-permissive temperature (26°C). White and black arrows indicate to the distal tip of the gonadal arm and vulva, respectively. Double headed arrow indicates to the expanse of a ‘nuclei mass’ located between ‘malformed’ oocyte and vulva. **(Biv)** Adult stage of an experimental dauer [*xrn-2*(*PHX25*)] after dauer-exit at non-permissive temperature (26°C). White and black arrows indicate to the distal tip of the gonadal arm and vulva, respectively. Double-headed arrow indicates to oocytes in ‘ectopic’ position. **(Bv, vi)** Adult stage of an experimental dauer [*xrn-2*(*PHX25*)] after dauer-exit at non-permissive temperature (26°C). Black and white arrows indicate to the distal tip of the gonadal arm and vulva, respectively. Double-headed arrow indicates to a region of ‘nuclei mass’ located between oocytes and vulva. Bvi depicts DAPI stained version of Bv. **(Ci)** Adult stage of a control dauer [*xrn-2*(*PHX25*)] after dauer-exit at permissive temperature (20°C). White arrow indicates to the distal tip of the gonadal arm. **(Cii, iii)** Adult stage of an experimental dauer [*xrn-2*(*PHX25*)] after dauer-exit at non-permissive temperature (26°C). Disoriented and ‘malformed’ oocytes are highlighted with double-headed arrow. Underlying solid line indicates to spermatheca harboring spermatids. Ciii depicts DAPI stained version of Cii. **(Di, ii)** Gravid Adult stage of an experimental dauer [*xrn-2*(*PHX25*)] after dauer-exit at non-permissive temperature (26°C). Dotted circles highlight oocyte nuclei. White double-headed arrow indicates to a region with ‘nuclei mass/ embryos’ that did not take up DAPI stain (Dii). **(E)** Short dauers [*xrn-2*(*PHX25*)] show reduced accumulation of indicated mature miRNAs at the non-permissive temperature during the early days (less than a week) after entering dauer state. **(F)** Reduced alteration in miRNA levels in short dauers is correlated with reduced larval arrests and germline anomalies after dauer-exit (compare with 7A left panel). **(G)** Continuation of perturbation of the endoribonuclease activity of XRN-2 post-dauer-exit (grown at 26°C) does not significantly contribute to the development of the germline anomalies observed in the short dauers [*xrn-2*(*PHX25*)] maintained/ grown at 26°C before and after dauer-exit.

Notably, at non-permissive temperature, *xrn-2*(*PHX25*) worms showed less dependence on the endoribonuclease activity of XRN-2 for determination of their mature miRNA levels, immediately after entering the dauer state (less than a week, to be referred as ‘short dauers’), in comparison to our ‘experimental dauers’ that were maintained for 30 days at 20°C before being subjected to the experimental condition (non-permissive temperature, 26°C; **Fig. 7E**). Interestingly, those short dauers subjected to the non-permissive temperature also displayed the same developmental defects upon dauer-exit as the ones displayed by the experimental dauers, albeit with diminished magnitude (**Fig. 7F**). It clearly suggested that miRNA metabolism during dauer state is not only critical for dauer metabolism, but also exerts effects, at least partly, on events that happen after dauer-exit, especially germline development and embryogenesis. It was reminiscent of an earlier observation, where it was suggested that transcripts synthesized and stored by pre-dauer worms facilitate dauer recovery as transcriptional inhibition of dauers could not prevent the initiation of that process^104^.

To understand whether the aforementioned defects in post-dauer worms (**Fig. 7F**) were only due to perturbation of the endoribonuclease activity of XRN-2 and associated deregulation of miRNAs that happened during the dauer stage or continuation of the nonpermissive 26°C temperature after dauer-exit, partly contributed to those phenotypes. Notably, *xrn-2*(*PHX25*) worms show defect in rRNA biogenesis in their continuous life cycle, when grown at the non-permissive temperature (26°C) for at least four generations^24^. Now, the short dauers (after subjected to 26°C for 110 hrs) were provided food and shifted to permissive temperature (20°C). Interestingly, we observed similar phenotypes with a penetrance level comparable to the short dauers that were maintained at 26°C after they were provided with food (**Fig. 7G**). These results clearly suggested that the endoribonuclease activity of XRN-2-mediated miRNA regulation is of paramount importance for the life cycle events after dauer-exit as the developmental features of the affected worms seemed to reflect loss of ‘molecular memory’ and traits of ‘precocious aging’, some of which had resemblance with the germline features observed in very old worms^94^. A thorough RNA sequencing-based analyses of these post-dauer-exit worms, especially of their germline transcriptome, would help us to understand the underlying molecular mechanisms for the anomalies observed, and how those are linked with the deregulated molecular pathways that operated during the dauer state.

## Discussion

Endoribonuclease activity of XRN-2 affected the mature levels of just over a couple of dozen miRNAs in dauers. However, it is possible that an increase in the sequencing depth might identify more miRNAs that are dependent on this activity. Notably, it is also possible that the paralogous protein, XRN-1, which is also known to act as a ‘miRNase’^105, 106^ might be playing a role in miRNA homeostasis in dauers in a much greater way to complement the role of XRN-2. This is a promising possibility as we have recently identified a highly efficacious endoribonuclease activity of this protein too (data not shown), which has so far been known to possess an exoribonuclease activity^107^. In the experimental dauers, one half of the affected miRNAs displayed accumulation of their mature forms compared to the control dauers without showing changes at the levels of their precursors (pri-and pre-miRNAs), with the only exception of *let-7*, which showed upregulation in transcription in the experimental dauers probably due to an increase in liganded DAF-12 activity (**Fig. 3A, D, E** and see results). But, as described before, in all likelihood, both increased abundance of pri-*let-7* and reduced turnover of the mature species contributed to the accumulation of its functional form. Conversely, some of the downregulated mature miRNAs displayed reduction in the levels of their primary transcripts. It could be possible that the TFs on which they are dependent for the synthesis of their pri-miRNAs were in turn downregulated due to the regulatory actions of some of the accumulated miRNAs. In support of this notion, we could find that transcription of *mir-232* and *mir-90* have been reported to get regulated by PRX-5, ODR-7, and NHR-34, FAX-1; respectively. The mRNA levels of these transcription factors showed downregulation in experimental dauers, and *prx-5 mRNA* and *nhr-34 mRNA* hosted target sites for *mir-253* and *lin-4,* respectively, which in turn were upregulated in the experimental dauers^108^. Overall, it was quite extraordinary to find that a small cohort of accumulated miRNAs that are dependent on the endoribonuclease activity of XRN-2 have predicted/ validated target sites in a large number of transcripts coding for components of pathways indispensable for the maintenance and survival of the dauers (**Fig. 1**). Importantly, many of those miRNA-target pairs (eg. *daf-12*/*Nr1h4* and *let-7/*let-7 sisters, daf-16/*Foxo6* and *let-7/*let-7 sisters, *daf-1*/*Sevr2a*/*2b* and *let-7/*let-7 sisters, *ilc-17.1*/ *Il17a* and *mir-80*/miR-450b etc.) are conserved and several of them have been linked to different disease states in humans, and thus of pathophysiological significance^109^. In this study, we have appreciated the ATP-independent endoribonuclease activity of XRN-2 in energy deprived dauers, considering it to be a constituent of the miRNasome-1 complex^23^, where it will remain active in the absence of inhibitory action by miRNasome-1.4, possibly due to its ATP-unbound state. Whereas, this endoribonuclease activity of miRNasome-1 would remain silent in ATP-replete continuously growing worms, at least in the L4/ young adult stages from where they were purified and studied. However, it is not inconceivable that in certain developmental stages in continuously growing worms (eg. in HSN neurons that remain quiescent undifferentiated until L4 stage), miRNasome-1 might bear modifications that might inhibit and alleviate the inhibitory action of ATP-bound miRNasome-1.4 on the endoribonuclease activity of XRN-2. Of course, one must not also exclude the possibility, where XRN-2 might be a member of a distinct complex, where its endoribonuclease activity could get exclusively used as ‘miRNase’ to shape the miRNA landscape.

In many ways, dauer quiescence is analogous to that of the quiescent state of the adult stem cells in higher eukaryotes and mammals. Dauer worms are heavily dependent on β-oxidation of fatty acids for their energy requirements and this entire pathway, at multiple steps, potentially gets regulated by miRNAs, whose accumulation in turn get modulated by XRN-2. This phenomenon has a strong connection with metabolic events that are critical for the homeostasis of several quiescent stem/ progenitor cells as well as cancer stem cells (CSCs)^110, 111^. It has been shown that hematopoietic stem cells (HSCs) rely on fatty acid oxidation (FAO) for stem cell maintenance and inhibition of FAO favors symmetric division and commitment, thus negatively affecting HSC maintenance and resulting in exhaustion of the HSC pool^110, 111^. ROS levels are also important for the activation of quiescent HSCs and correlate with HSC potency. Modulation of the balance between *de novo* lipogenesis (DNL) and FAO is also critical for fate determination (lymphoid vs myeloid) upon differentiation of HSCs. Lipid metabolism also plays an important role in the biology of neural stem/ progenitor cells (NSPCs)^110, 111^, and it is of significance not only during embryogenesis but also for neurogenesis in adulthood. Employing rodent models, adult neurogenesis has been studied in hippocampal dentate gyrus, subventricular zone of lateral ventricles (SVZ), hypothalamus, and has been shown to be critically important for learning and memory. Proliferating adult NSPCs upregulate the production of lipids, and genetic or pharmacological interference of this pathway is associated with a drastic reduction in proliferation and neurogenesis. The amount of DNL influences quiescence behavior as the production of new lipids is reduced in quiescent adult NSPCs. Importantly, FAO appears to be more important for quiescent NSPCs as they show higher levels of CPT1A-dependent FAO compared to proliferating NSPCs^110, 111^. Increased FAO was found to be high in adult NSPCs in mice SVZ and pharmacological inhibition of FAO resulted in reduced proliferation^110, 111^. Furthermore, a recent report linked the clinical association of FAO deficits with neuropsychiatric diseases to deregulation of NSPC activity during development^110^. Inhibition of FAO resulted in a reduced NSPC pool, due to increased differentiation and reduced self-renewal of NSPCs, suggesting that FAO is indeed crucial for the maintenance of NSPCs. In an Alzheimer mouse model, lipid accumulation led to decreased NSPC proliferation, which could be mimicked in wild type mice by a local increase and subsequent accumulation of lipids, suggesting that perturbed lipid metabolism during the disease state might be directly influencing NSPC behavior^110^. Together, there is now strong evidence pointing towards an important regulatory role of lipid metabolism in NSPCs. Similarly, FAO is also crucial for the maintenance of certain CSCs^110^. Mitochondrial FAO is initiated with the transfer of long-chain fatty acids from the cytosol into the mitochondrial matrix by carnitine palmitoyl transferase enzymes (CPT1 and CPT2). CPT1 is a rate-limiting enzyme in FAO, and is in the outer mitochondrial membrane, whereas CPT2 is located in the inner mitochondrial membrane. CPT1A is overexpressed in prostate cancer and is associated with a high tumor grade. CPT1B expression is enhanced in radiation-resistant breast cancer stem cells. High expression levels of CPT1A predict unfavorable clinical outcomes in AML and ovarian cancer. Genetic or pharmacological inhibition of CPT1A exerts anti-tumor activity in prostate cancer, melanoma, breast, and ovarian cancers^110, 111^. Increased expression of FAO linked genes is a hallmark in human GBM cells, and it is also the case in tumor-initiating stem-like cells (TICs) in HPCC. Similar to HSCs and NSPCs, high levels of FAO renders CSCs more quiescent-like, and such CSCs show high resistance to chemotherapy that targets proliferative cells^110^. Accordingly, upon inhibition of FAO, which is likely to make CSCs to proliferate, might make them more vulnerable to therapies. Now, mRNA levels of the components of CPT shuttle system such as *cpt-1, cpt-2, cpt-4, cpt-5, w03f9.4* got downregulated in the experimental dauers suggesting that FAO might have been diminished in them. Amongst those components, *cpt-1, cpt-2* and *w03f9.4* are predicted to have target sites for *mir-253*, *mir-77* and *mir-246*, respectively, and these miRNAs showed accumulation in the experimental dauers due to compromised endoribonuclease activity of XRN-2 (**Fig. 2A**). It is possible that hsXRN2 also performs a similar regulatory role, and indeed it has recently been implicated in miRNA turnover in humans as well^112^. Thus, hsXRN2 holds immense potential as a therapeutic target in modulating cellular fates, especially in post-injury/ degenerated tissues, which are dependent on metabolic processes linked to FAO.

In parasitic nematodes, infective larvae are also arrested, primarily as L3s^113^. They are non-feeding, survive harsh conditions, and analogous to *C. elegans* dauers. Thus, the molecular and cellular mechanisms used to control dauer decisions might have been co-opted to control infective larval metabolism and its recalibration upon finding host. Now, critical pathways involved in dauer maintenance did fail in the experimental dauers upon perturbation of the endoribonuclease activity of XRN-2 (**Fig. 3A-C)**. Additionally, stress chaperones such as *hsp-12.6*, *hsp-60* (members of *hsp-12.6* alpha-crystallin family) as well as ROS-response genes such as *sod-5, ctl-1, ctl-2, ctl-3* etc. also got downregulated in the experimental dauers (**Fig. 3B, C**), indicating that they might have lost their ability to sustain in a stressed condition. Moreover, genes involved in the catabolism of xenobiotic compounds were also downregulated in the experimental dauers. In a similar situation, in all likelihood, parasitic nematodes would have turned more susceptible to drugs (**Fig. 3B, C**). Therefore, the roles played by the endoribonuclease activity of XRN-2 in the infective lifecycle stages of parasitic nematodes need to be explored. Given the fact that there might be a species-specific positively charged RNA binding patch (PCP) critical for the endoribonucleolytic mechanism of XRN-2^24^, it holds potential for designing of effective drugs targeting parasitic nematodes affecting not only humans, but livestock and plants as well, and thus of relevance to animal husbandry industry and agriculture.

An interesting observation was that while *daf-2*, coding for the insulin-like receptor, positioned upstream in the IIS pathway was upregulated, but the PTEN ortholog DAF-18 was also upregulated (**Fig. 3A)**. DAF-18 promotes continuation of dauer state by inhibiting the suppression of DAF-16/FOXO through AKT signaling downstream of DAF-2 activation. But, still we observed that DAF-16 target genes were downregulated in the experimental dauers. As described before, this again indicated to the possible existence of a novel signaling pathway through which DAF-2/IIS pathway directly influence the function of DAF-16. Mutations in *PTEN*, a tumor suppressor gene, are often found to be the underlying pathological reason for the development and maintenance of various human cancers^114^. A human mutation that results in PTEN^C105Y^, which corresponds to DAF-18^C150Y^ in *C. elegans daf-18(yh1)* mutant, leads to an autosomal dominant syndrome, Bannayan–Riley–Ruvalcaba syndrome^47, 114^. This disorder is characterized by hamartomatous polyps in the intestine and benign subendothelial lipomas. Now, PTEN is considered to be of immense therapeutic importance in multiple human cancers. But, like the situation in worms, if as yet unknown parallel mechanisms would exist in PTEN-mediated pathways in humans too, then they might allow circumventing of the PTEN regulated wing of those pathways. Thus, a therapeutic intervention strategy designed only around PTEN might not achieve the desired outcome. Nevertheless, miRNA-mediated regulation of these conserved metabolic steps in worms and the underlying pivotal role played by XRN-2 to regulate those miRNAs appear to possess pathophysiological relevance.

We observed downregulated expression of multiple TFs, involved in terminal differentiation as well as maintenance of those differentiated neurons in experimental dauers, which probably happened due to the accumulation of different miRNAs that targeted the mRNAs of those candidates (**Fig. 4**). Notably, some of those terminal selector TFs (eg. UNC-30, UNC-86 etc) also play important roles in lineage specification during specific early developmental stages^59^. Thus, the experimental dauers provided us with a unique opportunity to study how simultaneous downregulation of different terminal selector genes results in the development of a complex neuronal problem, which is often the case in human neurodegenerative diseases^115^. Moreover, XRN-2-mediated regulation of those TFs at early developmental stages is of potential importance to understand the regulatory forces during early lineage commitments that are critical for the functional development of the nervous system.

Release of undesired cytokines as well as inappropriate quantities of *bona fide* cytokines often get referred as disequilibrated cytokine network, and when that happens above a critical level, it is called a cytokine storm^116^. It can interfere with normal cellular function and cause systemic alterations leading to heavy inflammation that can cause tissue damage and organ failure. Pathogenic infections, especially many viral infections often result in cytokine storms in patients. SARS-CoV-2 patients are known to display altered cytokine profiles that include changes in interleukins (IL-1, -6, -8, -17), TNFs and IFNs^116^. It has been suggested that one of the major factors contributing to cytokine storms in SARS-CoV-2 patients is increased biological activity of TGF-β that happens due to deregulation of cellular systems upon viral infection^117^. TGF-β has a broad spectrum of biological activities in humans and acts as an endogenous pyrogen. Deregulated levels of it can lead to fever, mitochondrial dysfunction, and affect Na^+^-K^+^-ATPase activity, and these last two events cause fatigue, which is a typical symptom of SARS-CoV-2 patients. Dry cough and changes in lung interstitium are also caused by increased TGF-β by means of activating fibroblast growth factors (FGFs). High TGF-β also promotes heavy secretion by bronchial mucus cells that hinders normal breathing, and can lead to serious infections and shock. Additionally, TGF-β is a strong immunosuppressive factor in the body, inhibits proliferation and differentiation of lymphocytes, and thus suppresses immune function and causes delay in recovery. TGF-β negatively impacts olfactory and gustatory receptor neurons and neurogenesis by inhibiting progenitor cell proliferation and differentiation, and its deregulation gets manifested in the loss of olfactory and gustatory senses in some patients^117, 118^. Overall, the clinical manifestations of SARS-CoV-2 are highly consistent with increased TGF-β activity, which is thought to be one of the main reasons for occurrence of cytokine storms. Here, in the experimental dauers, we have reported downregulation of mRNAs coding for DAF-7 (TGF-β) and one of its receptor subunits (DAF-4), as a likely consequence of heavy accumulation of targeting miRNAs (**Fig. 3A**). Moreover, ILC-17.1 (IL-17) as well as its receptor got downregulated at the mRNA levels in the experimental dauers, potentially for the same reason (**Fig. 5A**). Elevated IL-17 levels have already been reported to facilitate cytokine storms and have been linked with viral load and disease severity. Interestingly, elevated IL-17 levels were also previously described in patients with MERS-CoV^119^, and thus appears to be a common pathological signature in patients infected with this particular family of viruses. Therefore, XRN-2 appears to possess a potential regulatory role in this immuno-inflammatory-axis and needs to be carefully assessed for its therapeutic potential.

XRN-2 is ubiquitously expressed, and XRN-2 regulated miRNAs appear to play important roles in modulating the expression of critical components of several pathways central to the biology of multiple organs/ systems, and thus proved to be indispensable for the survival of quiescent dauers (**Fig. 1A, Fig. 8**). Most of the revelations on miRNA metabolism in worms have been proven to be conserved^109, 120^, and the recent finding that the human homolog of worm XRN-2 regulates mature miRNAs that are of pathophysiological significance in several human cancers further strengthened this notion^112^. Therefore, it does not require any stretch of imagination to postulate that hsXRN2 might be playing regulatory roles in similar/ identical pathways, as described in this study, which are operational in equivalent cellular states and beyond, and thus it might be of paramount importance to human physiology, both in health and disease. Finally, pathogenic molecules/ peptides, including viral peptides, are known to target proteins and enzymes central to human metabolism, a strategy adopted to debilitate/ hijack the host system^121^. Accordingly, it was shown that the SARS-CoV-2 spike protein interferes with key DNA damage repair protein BRCA1, as well as with 53BP1, and thereby potentially affects V(D)J recombination process central to adaptive immunity^122^. hsXRN2, a key player in cellular homeostasis, in all likelihood, might also be a potent target of pathogenic and viral peptides/ molecules. Thus, molecular docking studies involving an atomic structure of hsXRN2 might facilitate the first step to appreciate the possible pathophysiological consequences of interference of its function upon infection, and that will not only augment vivid understanding of the therapeutic potential of hsXRN2 but might also pave way for designing of strategies for clinical interventions.

**Fig. 8.**
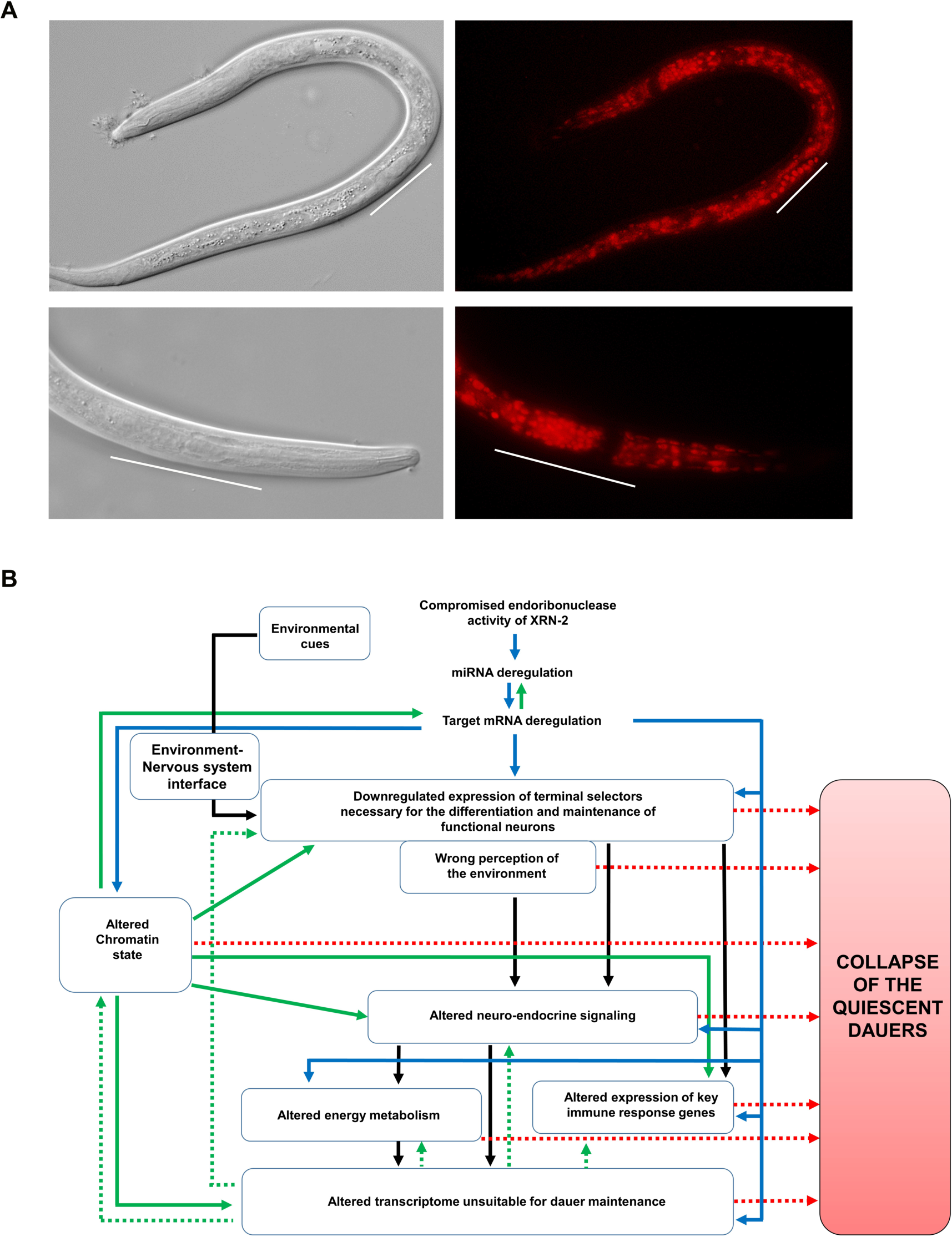
**(A) XRN-2 (*xrn-2::mCherry*) shows widespread expression in dauers.** Representative epifluorescence images on the right and the corresponding DIC images on the left. **Top panels.** White bars indicate position of the gonad in a dauer worm. **Bottom panels.** White bars indicate approximate pharyngeal region in a dauer. **(B) A putative model depicting importance of the endoribonuclease activity of XRN-2-mediated regulation of miRNAs contributing to homeostasis and integration of metabolism and functions in and between multiple systems in dauers.** Solid blue arrows indicate to putative direct effects. Solid black arrows indicate to putative direct and/or reinforcing effects. Solid green arrows indicate to putative secondary effects. Dotted green arrows indicate to putative tertiary effects. Dotted red arrows indicate to putative factors that may contribute to the collapse of the sturdy quiescent dauers.

## Author Contributions

SC designed the research and experiments. TC, AS and MS performed the experiments. Together, all the authors analyzed the results and SC wrote the manuscript.

## Acknowledgments

We sincerely thank Rafal Ciosk for his insights on worm germline anomalies reported in this study. This work is not supported by any funding agency/ organization. TC, AS, MS received fellowships from KVPY-MHRD (Govt. of India), CSIR (Govt. of India), and UGC (Govt. of India), respectively.

