## Supplementary Information for "Dauer quiescence as well as continuity of the life cycle after dauer-exit in *Caenorhabditis elegans* are dependent on the endoribonuclease activity of XRN-2"

**Laboratory of RNA Molecular Biology**

**Indian Institute of Science**

**Bangalore, India**

**#** Contributed equally

**This file includes:**

Materials and Methods

**Nematode Strains and Maintenance.** *C. elegans* were grown under standard conditions using *Escherichia coli* strain OP50 as food source**^123^**. Bleaching followed by starvation-induced L1 arrest was used to generate synchronized cultures. The strains used in this study are *N2* and *xrn-2(PHX25)*.

**XRN-2 inactivation in dauer worms.** *xrn-2(PHX25)* worms were pushed into dauer stage through starvation and over-crowding. Then randomly ten worms were picked from each plate for microscopic confirmation for the presence of mouth-plug and atrophied pharynx, which are distinctive features of dauer worms. Plates from where at least seven out of ten worms tested positive for the aforementioned dauer phenotypes were selected, and further maintained for four more weeks. To perturb XRN-2’s endoribonucleolytic activity, half of those starved dauer worm plates were shifted to the non-permissive temperature of 26ºC (experimental dauers), whereas the other half was maintained at the permissive temperature of 20ºC (control dauers). Dead worms were counted at 24 hr intervals, and removed with a platinum wire. Survival analysis (n = 1000) was performed using Kaplan-Meier estimate, and the survival curve was plotted using GraphPad Prism. After 110 hrs of dwelling under the two different conditions (permissive vs non-permissive temperature), the control and experimental dauer worms were harvested and subjected to 1% SDS treatment for 30 minutes to eliminate any non-dauer worm, and further subjected to several molecular analyses.

**RNA isolation, Northern blotting and RT-qPCR analysis.** For total RNA isolation, animals were harvested at young-adult stage from plates by washing with M9 buffer (22 mM KH_2_PO_4_, 22 mM Na_2_HPO_4_, 85 mM NaCl, 1mM MgSO_4_). Animals were pelleted and suspended in ten pellet volumes of TRIzol reagent (Ambion), and total RNA was extracted according to the manufacturer’s protocol. Northern blotting of endogenous RNA was done as described**^124^**. In short, for detecting small RNA, 30 μg of total RNA was resolved on 7 M urea/ 12% PAGE, transferred to Hybond N+ (GE Healthcare) membrane and probed appropriately. Hybridization was carried out overnight using 5’-end labeled DNA probe.

For RT-qPCR, 1 μg of total RNA of the respective samples were reverse transcribed using SuperScript™ III Reverse Transcriptase (Invitrogen) in 20 μl reaction containing 5 mM DTT, 0.5 mM dNTPs, 2.5 μM oligo (dT)_20_ and 1.0 μl of enzyme, in accordance with the manufacturer’s protocol. 1.5 μl of each of the reverse-transcription reactions were amplified with PowerUp^TM^ SYBR Green Master Mix (Applied Biosystems) in a total volume of 20 μl using specific forward and reverse primers at a concentration of 0.5 μM each, and analyzed in a ViiA™ 7 Real-Time PCR System using ΔΔCt method. Values were normalized against the value of *eft-2* expression. The experimental values, representing means (±SEM) from three independent biological replicates, were compared to the expression levels of the corresponding control samples. TaqMan MicroRNA Assays (Applied Biosystems) to quantify individual miRNAs were performed as per the supplier’s protocol and as described **^118^**. In brief, 10 ng of total RNA of the respective samples, were reverse transcribed in 7.5 μl reaction volume and then 0.67 μl of each of the reverse-transcription reactions was amplified in a total volume of 20 μl, and analyzed in a ViiA™ 7 Real-Time PCR System using ΔΔCt method. Values were normalized against the value of *sn2841* and *U18* expression in the respective samples.

### Small RNA cloning. Total RNA was extracted from control and experimental dauers using TRIzol reagent (Ambion). Small RNA was enriched using MirVana Kit (Ambion). RNAs ranging from 18–40 nt were further purified employing 7 M urea/ 15% PAGE. One μg each of purified small RNA samples were treated with RNA 5’- Pyrophosphohydrolase (NEB), which allows for cloning of 5’-capped/ triphosphate carrying small RNAs, at 37°C for one hour. The reactions were stopped with the addition of phenol:chloroform, and the treated small RNAs were purified via ethanol precipitation. Sequencing libraries were prepared using the NEBNext Multiplex Small RNA Library Prep Set for Illumina (NEB). PCR amplicons were size selected on 10% polyacrylamide gels. The libraries were sequenced on Illumina HiSeq platform.

**mRNA Cloning.** DNase-treated total RNA with RIN > 8 was used to prepare strand-specific RNA libraries. Firstly, poly(A)+ RNA was isolated from total RNA using NEBNext® Poly(A) mRNA Magnetic Isolation Module (NEB). Then strand-specific RNA-seq libraries were prepared using the NEBNext® Ultra™ II Directional RNA Library Prep Kit (NEB) according to the manufacturer’s instructions.

### Oil Red O staining and analysis.  Oil Red O (ORO) staining was performed as mentioned earlier^125^.

### mRNA sequence analysis. Reads were adapter-trimmed using fastp v0.20.0^126^ (fastp --in1 "$sample_R1.fq.gz" --in2 "$sample_R2.fq.gz" --out1 "$sample_R1.trim.fq.gz" --out2 "$sample_R2.trim.fq.gz" -q 20 -u 10 --detect_adapter_for_pe –correction). Trimmed reads were then aligned to the *C. elegans* genome (genome release WS280 from WormBase^127^) using the HISAT2^128^ splice-aware aligner with default settings. The samtools^129^ collate-fixmate-markdup pipeline was used to eliminate optical duplicates in the resulting .bam file, and reads were assigned to features in the canonical geneset using Subread featureCounts 2.0.3^130^ (featureCounts -a ws280.canonical_geneset.gtf -p --countReadPairs -C -M -O --fracOverlap 0.01 --fraction --ignoreDup --largestOverlap -o "$output" "${aligned_files[@]}"). The fractional read counts output from featureCounts was then analyzed using DESeq2^131^ to identify differentially expressed genes (DEGs), defined as genes with $\boldsymbol{FC}\boldsymbol{\leq}\boldsymbol{0}\boldsymbol{.}\boldsymbol{7} \text{or} \boldsymbol{FC}\boldsymbol{\geq}\boldsymbol{1}\boldsymbol{.}\boldsymbol{3}$, and $\boldsymbol{p}_{\boldsymbol{adj}}\boldsymbol{\leq}\boldsymbol{0}\boldsymbol{.}\boldsymbol{1}$.

### small RNA sequence analysis. Reads were adapter-trimmed using fastp v0.20.0^126^ (fastp -w 8 -i ${sample}.fq.gz -o ${sample}.trim.fq.gz --length_required 16) using automatic adapter detection. Reads smaller than 16 nt after adapter and quality trimming were discarded. The remaining reads were converted from FASTQ to FASTA and subjected to the miRdeep2-quant^132^ pipeline, with the miRbase v22 release as reference (mapper.pl "${sample}.trim.fq" -e -m -l 16 -h -s "${sample}.collapsed.fa" -v; quantifier.pl -d -j -W -p "$ref_hairpin" -m "$ref_mature" -P -r "${sample}.collapsed.fa" -y "${sample}";). The resulting count files for each sample were combined into one CSV file using a Python script. These counts were then normalized to the external spike-in has-miR-122-5p, which was assigned a read of 10,000 in each sample to adjust for variable sequencing depth. These normalized counts were then analyzed by DESeq2^131^ to identify differentially expressed miRNAs, which were defined as miRNAs possessing $\boldsymbol{FC}\boldsymbol{\leq}\boldsymbol{0}\boldsymbol{.}\boldsymbol{7} \text{or} \boldsymbol{FC}\boldsymbol{\geq}\boldsymbol{1}\boldsymbol{.}\boldsymbol{3}$, and $\boldsymbol{p}_{\boldsymbol{adj}}\boldsymbol{\leq}\boldsymbol{0}\boldsymbol{.}\boldsymbol{1}$.

### mRNA-miRNA correlation analysis. miRNAs targeting the 3’ UTR of DEGs were predicted using the TargetScanWorm^130^, release 6.2 database. Briefly, all miRNAs possessing a seed sequence known to target the UTR of a known transcript were assigned as miRNAs targeting the gene. The geometric mean of the fold changes of the differentially-expressed miRNAs (identified in the previous step) targeting each DEG was calculated, and contrasted with the fold change of that DEG.

### GO enrichment and pathway enrichment analysis. Gene ontology term enrichment analysis of DEGs and DAF-16/DAF-12 target genes was performed using the g:OSt tool from the g:Profiler web service (<https://biit.cs.ut.ee/gprofiler/gost>), version e105_eg52_p16_e84549f^26^, using the Gene Ontology: Biological Process^134^, KEGG^135^ and WikiPathways^136^ databases, with all other settings at their recommended defaults for *C. elegans*. Non-redundant GO terms were manually curated, and plots were generated using a custom Python script. All custom code related to this project is at <https://github.com/tamchow/mirnasome_1/tree/main/sc25>.
